# Evolution of CTCF underlies the establishment of loop extrusion as a predominant chromatin folding mechanism in vertebrates

**DOI:** 10.64898/2026.10.19.759930

**Authors:** Liene Astica, Rafael D. Acemel, Henrike Schwarzenbacher, Claudia Racioppi, Ana M. Rojas, Josep Rotllant, Ozren Bogdanovic, Francisca M. Real, Lionel Christiaen, Robert Zinzen, Darío G. Lupiáñez

**Affiliations:** Max-Delbrück Center for Molecular Medicine in the Helmholtz Association (MDC), Berlin Institute for Medical Systems Biology (BIMSB), Epigenetics and Sex Development Group, Berlin, Germany; Max-Delbrück Center for Molecular Medicine in the Helmholtz Association (MDC), Berlin Institute for Medical Systems Biology (BIMSB), Systems Biology of Neural Tissue Differentiation Group, Berlin, Germany; Centro Andaluz de Biología del Desarrollo (CABD), CSIC/UPO/JA, Seville, Spain; Humboldt-Universität zu Berlin, Faculty of Biology, Berlin, Germany; Department of Biology, New York University, New York, NY, USA; Regeneron Pharmaceuticals Inc, Tarrytown, NY, USA; Aquatic Biotechnology Lab., Institute of Marine Research, Spanish National Research Council (IIMCSIC), 36208, Vigo, Spain; Michael Sars Centre, University of Bergen, Bergen, Norway; Max-Delbrück Center for Molecular Medicine in the Helmholtz Association (MDC), Advanced Light Microscopy Technology Platform (ALM), Berlin, Germany

## Abstract

CTCF is a DNA-binding protein that emerged in the last bilaterian ancestor and that plays a central role in 3D genome organization. Yet, despite its relevance, genomic studies have revealed notable differences in CTCF function across animal clades. In vertebrates, CTCF is essential for TAD organization through cohesin-mediated loop extrusion, whereas its contribution to chromatin organization outside this clade appears to be more limited. Here we investigate the evolution of CTCF as an architectural factor through a cross-species functional replacement approach, in which orthologs from divergent bilaterian lineages are expressed in either mouse embryonic stem cells or *Drosophila* embryos. Chromosome conformation analyses demonstrated that the ability of CTCF to mediate loop extrusion-dependent TAD formation is encoded within the vertebrate protein. Yet, this property alone is insufficient to establish vertebrate-like genome architecture in an invertebrate background, suggesting the existence of additional lineage-specific factors. ChIP-seq analyses further revealed that CTCF DNA-binding specificity has been progressively refined during bilaterian evolution, with the vertebrate lineage exhibiting a preferential shift towards distal genomic regions that is consistent with an expansion of regulatory domains. Notably, despite failing to form TADs, the ortholog from a non-vertebrate chordate partially rescued the lethality associated with the loss of the vertebrate CTCF protein, indicating that its architectural and regulatory functions are evolutionary independent. Further Hi-C analyses in tunicate and cyclostome species suggest that the ability of CTCF to act as a loop extrusion barrier was already present in the last common ancestor of all vertebrates. Altogether, our findings establish the diversification of CTCF regulation and function as key drivers for the specialization of bilaterian 3D genome organization.

## INTRODUCTION

Animal genomes are organized into three-dimensional (3D) chromatin domains that shape the functional interactions between regulatory elements and their target genes^1,2^. Genomic rearrangements that alter 3D chromatin domains can cause human disease, including developmental malformations or cancer^3–6^. Conversely, analogous genomic changes can also provide a powerful substrate for evolutionary adaptation by rewiring regulatory interactions across such domains^7–10^. Despite the biological importance of 3D chromatin domains, studies across diverse animal models have revealed remarkable differences in the mechanisms underlying their formation^11^. In vertebrates, 3D genome organization emerges from the interplay between two distinct mechanisms^12^. First, the segregation of megabase-sized genomic regions according to epigenetic status originates spatially distinct A/B compartments. Second, cohesin-mediated loop extrusion drives the formation of Topologically Associating Domains (TADs), which can span hundreds of kilobases (kb) and reflect the existence of regulatory landscapes^13–17^. This influential framework, first established in mammals, has been subsequently extended to other vertebrate groups, including birds^18^, amphibia^19^, bony fishes^20^ and cartilaginous fishes^9^. In invertebrate species, early chromatin conformation studies also identified chromatin domains that were reminiscent of vertebrate TADs^21^. Yet increased experimental resolutions, simulation approaches and functional perturbations have provided a more nuanced view. In *Drosophila*, the most extensively studied invertebrate model, 3D genome organization is predominantly driven by transcriptional activity and epigenetic status^22^, whereas loop extrusion is mainly associated with enhancer-promoter interactions rather than chromatin domain formation^23^. Similar signatures have been reported for other invertebrate lineages, including echinoderms^24^. Collectively, these observations have created an important conceptual conundrum in the field, as chromatin domains displaying similar visual patterns in experimental data are recurrently classified under the same terminology despite arising through distinct molecular mechanisms^1,25^. More broadly, these findings raise questions about how the mechanisms governing 3D chromatin organization have evolved and how they ultimately shape developmental processes.

The insulator protein CCCTC-binding factor (CTCF) is a multifunctional protein that emerged in the bilaterian ancestor^26–29^. CTCF binds to DNA through a conserved 11 zinc finger (ZF) domain, with ZF3-7 being the crucial residues for motif recognition^30–32^. Besides its functions in transcriptional regulation or cell division, CTCF has gained particular attention for its central role in vertebrate genome organization as a modulator of cohesin-mediated loop extrusion. During interphase, the cohesin complex extrudes the chromatin fiber, bringing distant genomic regions into close spatial proximity^33^. Cohesin is preferentially stabilized at genomic sites where CTCF is bound to its cognate DNA motif in a convergent orientation, exposing the N-terminal domain of the protein to the approaching complex^34,35^. Specific aminoacidic sequences within the vertebrate CTCF N-terminus have been implicated in cohesin stabilization, including the YXF and KTYQR motifs^36–38^. Consequently, pairs of distal CTCF sites can interact with each other, defining the boundaries of TADs and originating a characteristic focal enrichment signal at the apex of these domains that is observable in Hi-C maps. Given its multifunctional nature, the constitutive depletion of CTCF impairs cell viability or normal development^20,39–41^, causing defects in chromosome segregation and cellular division. Thus, degron strategies for acute degradation were developed to elucidate its precise role as modulator of 3D chromatin organization. The rapid depletion of CTCF^15^, similarly as it occurs for the cohesin complex^16^, causes the disappearance of TADs, and the preservation or even reinforcement of A/B compartment signal. Given the relative conservation of both CTCF protein structure and of its DNA-binding motif, analogous architectural functions were hypothesized to exist across the Bilaterian clade. Surprisingly, CTCF was shown to be largely dispensable for early development in *Drosophila*, as mutant fly embryos progress until adulthood only displaying restricted phenotypes associated to *Hox* gene dysregulation^42^. Consistent with these observations, Hi-C analyses revealed that 3D genome organization is largely preserved in the absence of CTCF, with loss of insulation in a subset of boundary regions^43^. Moreover, the impairment of loop extrusion in *Drosophila* only revealed specific changes in enhancer-promoter interaction, while chromatin domains remained largely unaffected^23^. These results, together with observations from other invertebrate species, suggest that loop extrusion may not be the predominant mechanism underlying chromatin folding outside vertebrates. In this context, the role that CTCF could have played in the emergence of distinct modes of 3D chromatin organization during bilaterian evolution remains unclear.

Here we investigated the evolution of CTCF as a genome architectural factor by performing orthologous replacements from representative species of distinct bilaterian lineages, in both mouse embryonic stem cells (mESC) and *Drosophila* embryos. We exploit this context-controlled system to address how these distinct CTCF homologs influence 3D chromatin organization, transcriptional regulation, and organismal phenotypes. Through this setup, we demonstrate that the CTCF protein has conserved its function as a barrier for loop extrusion across vertebrate evolution, being necessary but not sufficient to mediate the formation of TADs. Furthermore, we show that CTCF can operate as a transcriptional regulator and as an architectural factor, and that both functions are independent and evolutionary uncoupled. By extending chromatin conformation analyses to key lineages, we trace the emergence of CTCF function as a barrier to loop extrusion, confirming that it precedes the diversification of vertebrates. Collectively, our results delineate a model in which CTCF has played a major role in the evolution of 3D chromatin organization through shifts in chromatin-binding dynamics and functions.

## RESULTS

### *The capacity of CTCF to form TADs via loop extrusion is encoded in the vertebrate* protein

Since CTCF is widely present across bilaterian species^26^, we investigated whether orthologs from different clades are functionally interchangeable. As CTCF inactivation is lethal in mESCs, we developed a system allowing the acute degradation of the endogenous mouse protein while maintaining the expression of a species-specific ortholog. Briefly, we engineered mESCs to carry an auxin-inducible degron system that enables the rapid depletion of the endogenous CTCF (**Figure 1A**), an approach that has been widely used to study the function of this factor^15^. As expected, the treatment with indole-3-acetic acid (IAA) resulted in a complete degradation of the CTCF protein (**Figure 1B**). We further modified these cells by replacing one of the degron-tagged alleles with the *CTCF* coding sequence (CDS) from a different species. Specifically, an orthologous *CTCF* CDS, fused to the mouse 3′UTR, was inserted immediately downstream of the endogenous start codon (ATG) through CRISPR/Cas9-mediated homologous recombination. This modification maintains the endogenous regulatory configuration of the mouse *Ctcf* locus, including the promoter, 5′ and 3′ UTRs, and the first intron, ensuring that any observed effects are attributable solely to differences in CDS. The replaced genes were engineered to encode a FLAG-tag, enabling protein immunoprecipitation for all species orthologs using a common antibody. Upon IAA treatment, the endogenous mouse CTCF is depleted from one allele, leaving only the expression of an orthologous protein from the other one.

**Fig. 1.**
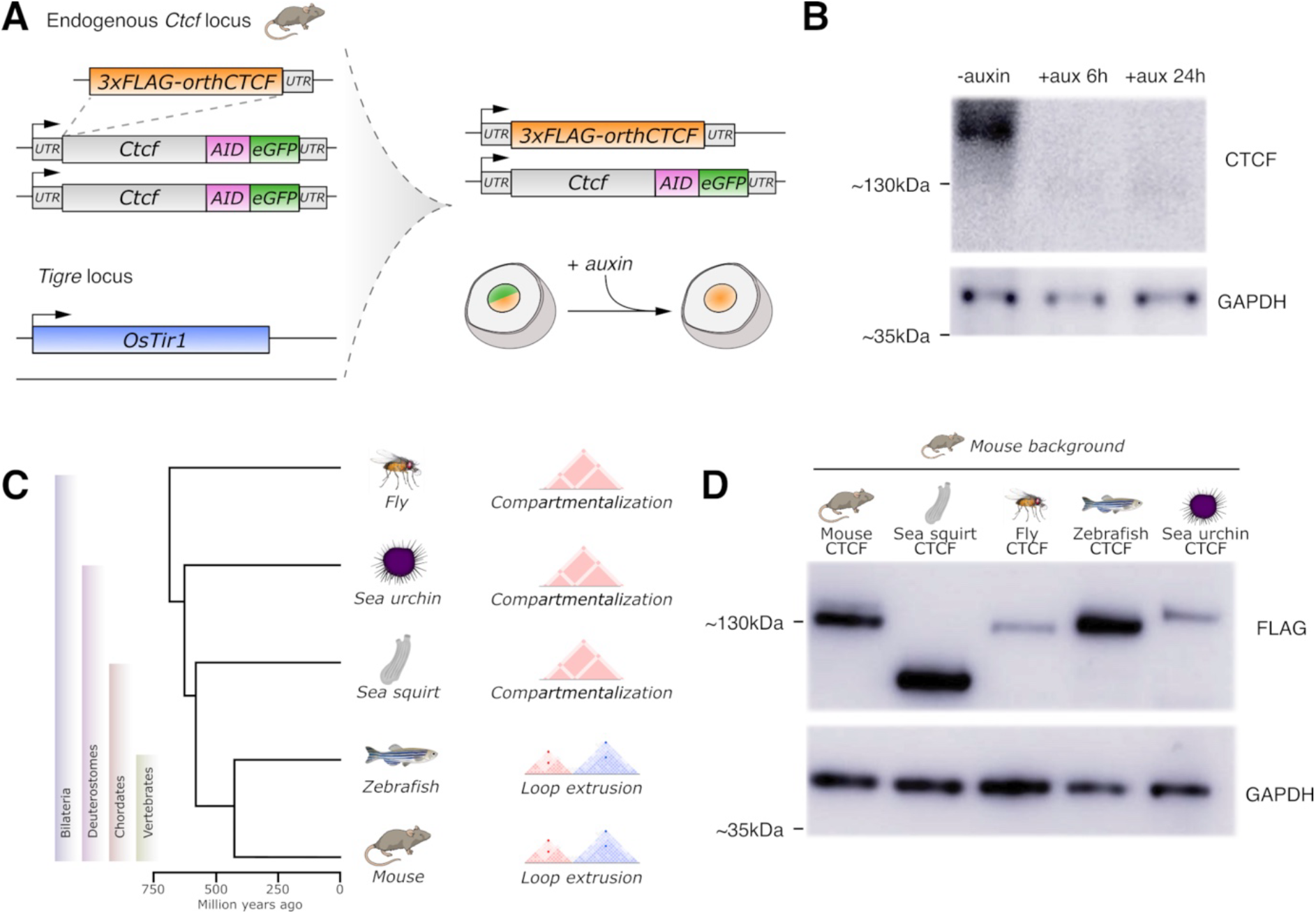
CTCF orthologous replacement in mESC. **A.** Experimental strategy to generate mutant mESC carrying orthologous CTCF replacements. Initially, mESC were modified to integrate an AID-tag followed by a GFP marker downstream of the endogenous Ctcf, in homozygosity, as well as the OsTir1 gene at the Tigre locus. One of the Ctcf alleles was further modified to introduce a FLAG-tagged CDS version of CTCF from a selected species. This configuration allows the constitutive expression of the orthologous gene. Upon IAA treatment, the system allows the rapid depletion of the endogenous CTCF protein, while the orthologous version remains to be expressed. **B.** Western blot analyses on mESC carrying auxin-inducible degron system. Note the absence of detectable CTCF protein upon addition of IAA, after 6h and 24 h. **C**. Selection of representative species for relevant bilaterian clades. Left, evolutionary tree showing divergence time between species. Right, predominant mechanisms associated with the formation of chromatin domains. **D.** Western blot analysis using a common anti-FLAG antibody. Note the presence of a specific band with the expected sizes for all CTCF orthologs in each of the mESC replacement lines.

To explore the evolution of CTCF as a 3D chromatin organization factor, we performed orthologous replacements using CDS from species that are strategically positioned across the animal phylogeny (**Figure 1C**). First, we generated a replacement using the CTCF CDS from the zebrafish *Danio rerio*, a vertebrate species that diverged from mouse approximately 400 million years ago and where TADs have been demonstrated to form via loop extrusion^20^. We also performed a replacement using the CTCF CDS from the sea squirt *Ciona robusta*, a non-vertebrate chordate that separated from the mouse approximately 550 million years ago. In this species, we reanalyzed an existing Hi-C dataset on tailbud embryos^44^ and, although we could detect large-scale chromatin compartments, we found no evidence of TADs formation (**Supplementary Figure 1**). Since cellular heterogeneity may potentially mask TAD signals, we performed analogous experiments in FACS-sorted heart and mesenchymal cells, further confirming the absence of signals associated with loop extrusion-mediated domain formation (**Supplementary Figure 1**). We also generated a replacement line with the CTCF CDS from the sea urchin *Strongylocentrotus purpuratus*, a deuterostome species that diverged more than 600 million years ago, and where chromatin domain formation also appears to be largely independent of loop extrusion^24^. Finally, we included the *Drosophila melanogaster* CTCF ortholog, representing an evolutionarily distant bilaterian lineage that diverged more than 700 million years ago. Notably, fly CTCF has been demonstrated to have distinct roles in 3D genome organization and embryonic viability compared to mouse^42,43^. For all the studied species, the CTCF CDS is well annotated and contain 9-11 Zinc Fingers (ZF), except for the sea squirt. In this species, the gene model annotated in the Aniseed database^45^ only contained 8 ZF and lacked any C-terminal portion. To verify the accuracy of this annotation, we performed 3′RACE on *Ciona robusta* embryonic tissue, confirming the reduced number of ZF and retrieving its C-terminal region. Therefore, despite extensive evolutionary divergence across these bilaterian clades, CTCF has largely conserved ZFs 3–7, which mediate binding to DNA^31,32^ (**Supplementary Figure 2**). In addition, the N-terminal domain of the protein of all species contains a YXF residue that is critical for interaction with the cohesin complex^36,37^ (**Supplementary Figure 2**). As a technical control for the orthologous replacement lines, we generated an additional one carrying the mouse CTCF CDS. In all cases, the engineered mESCs expressed the corresponding orthologous protein and, upon IAA treatment, the endogenous mouse CTCF was efficiently degraded (**Figure 1D**).

To evaluate the effects of CTCF orthologous replacement on 3D chromatin organization, we generated Hi-C maps for each mESC replacement line after acute degradation of the endogenous CTCF. We first assessed chromatin compartmentalization by calculating the first eigenvector (PC1). Across all CTCF replacement conditions, A and B compartments were overall preserved (**Supplementary Figure 3**). These effects were analogous to those observed upon complete CTCF degradation, indicating that chromatin segregation by homotypic interactions is largely independent of CTCF and not driven by the species-specific properties of the orthologs. We next inferred the presence of TADs, by calculating boundary scores. In all mESC lines expressing invertebrate CTCF orthologs (fly, sea squirt and sea urchin), TAD structures were absent (**Figure 2A** and **Supplementary Figure 4**). Moreover, the interaction profiles of lines expressing the invertebrate CTCF orthologous were similar to those from degron cells treated with IAA, that is with complete degradation of the endogenous protein (**Supplementary Figure 4**). These results demonstrate that invertebrate CTCF orthologs are not sufficient to mediate TAD formation via loop extrusion in an otherwise vertebrate context. In contrast, replacement lines expressing the zebrafish CTCF displayed a complete preservation of TAD organization (**Supplementary Figure 4**). Hi-C maps displayed the characteristic focal interaction peaks at TAD corners, an indicative of loop extrusion–driven organization. Furthermore, the interaction profiles from these cells were virtually indistinguishable from mouse wildtype or degron controls without IAA (**Supplementary Figure 4**), demonstrating that vertebrate orthologs can effectively replace the function of the endogenous mouse CTCF protein. The calculation of boundary scores across all replacement and control lines revealed two distinct distributions: one comprising all invertebrate replacements as well as the CTCF degron line treated with IAA, and another including zebrafish and mouse orthologs, together with untreated and wildtype control lines (**Figure 2B** and **Supplementary Figure 5**). Altogether, these results indicate that the ability of CTCF to mediate TAD formation through loop extrusion is encoded within the vertebrate protein and has been preserved despite the ∼400 million years of evolutionary divergence that separate bony fishes and mammals.

**Fig. 2.**
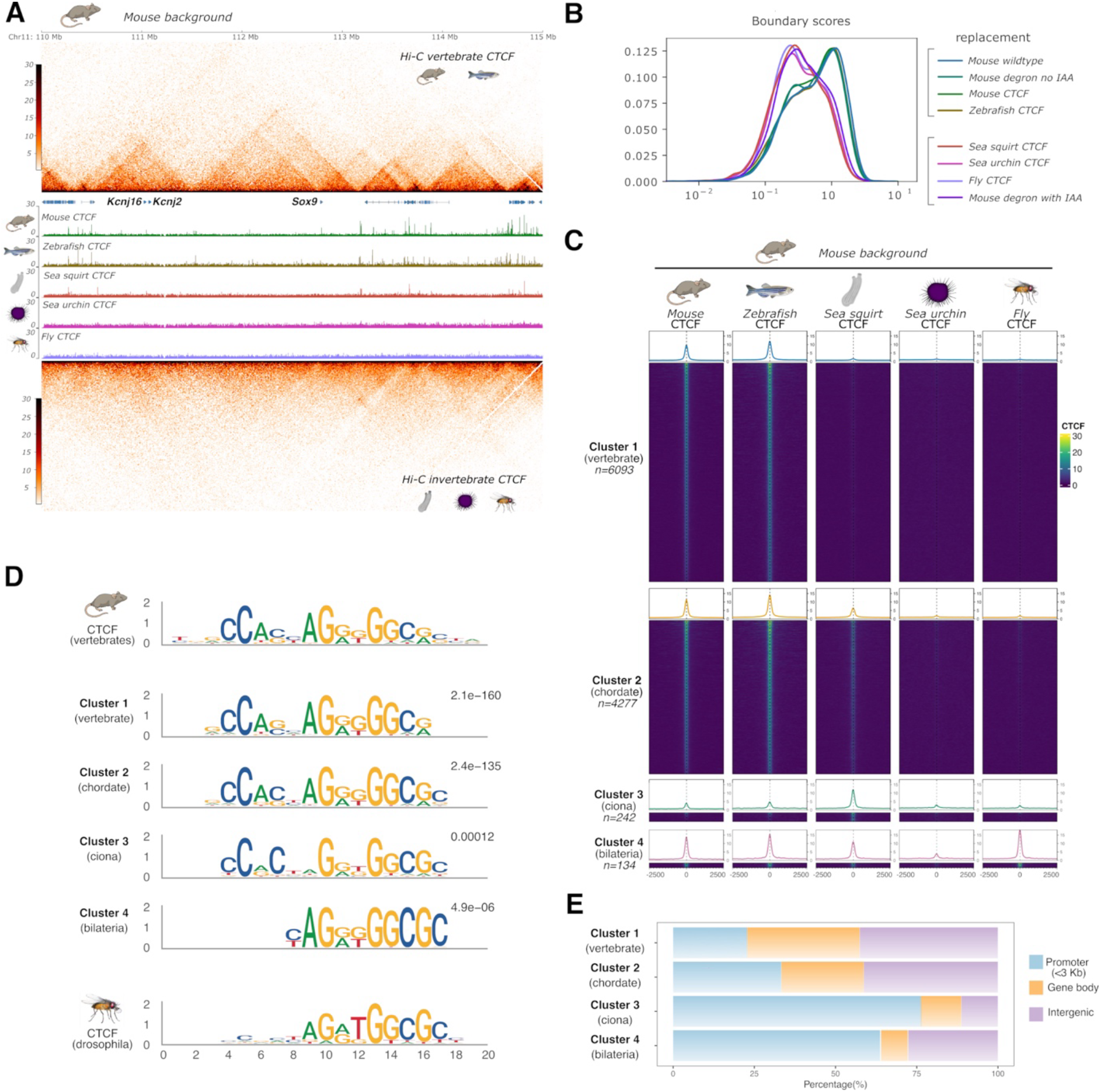
Functional effects of CTCF orthologous replacement in mESC. **A.** Effects of CTCF replacement on vertebrate chromatin in a representative genomic region (Sox9 locus). Top and bottom, Hi-C maps from vertebrate and invertebrate replacements. Displayed Hi-C maps correspond to zebrafish for vertebrate and sea squirt for invertebrates, but orthologous replacements from the other species show analogous results (see Supplementary Figure 4). Middle, ChIP-seq profiles of CTCF ortholog binding. Note the lack of TADs in sea squirt replacement, despite binding of CTCF onto chromatin. All ChIP-seq experiments were performed using a common antiFLAG antibody **B.** Distribution of boundary scores from Hi-C experiments. Note the presence of two distribution regimes, containing invertebrate and vertebrate samples, respectively (brackets). **C.** Clustering of CTCF ChIP-seq signal at specific genomic regions. **D.** Most represented motif per ChIP-seq cluster. The canonical CTCF motif described for the vertebrates and fly is included for reference. **E.** Genomic distribution of CTCF binding events per each ChIP-seq cluster.

### *Invertebrate CTCF orthologs bind across vertebrate genomes without altering* chromatin folding

Given the lack of TAD organization in mESC lines expressing invertebrate CTCF orthologs, we next assessed the ability of these proteins to bind the vertebrate genome. To this end, we took advantage of the FLAG-tag incorporated into each orthologous replacement to perform ChIP-seq experiments using a common anti-FLAG antibody, enabling a direct comparison of experiments across all lines (**Figure 1A**). Importantly, the insertion of the FLAG-tag did not produce detectable effects on 3D chromatin organization or CTCF genome-wide binding, as confirmed in mouse control cells (**Supplementary Figure 4**). The inspection of ChIP-seq profiles revealed that mouse and zebrafish mESC lines displayed nearly identical CTCF binding patterns, indicating that vertebrate orthologs recognize and bind to the same genomic regions (**Figure 2A**). Intriguingly, the sea squirt ortholog was often bound to sites normally occupied by the vertebrate CTCF, despite its inability to mediate TAD formation (**Figure 2A**). Sea urchin and fly CTCF proteins likewise exhibited detectable binding across the vertebrate genome, although at lower levels than those observed for vertebrate orthologs (**Figure 2A**). Together, these results demonstrate that, despite their evolutionary divergence, CTCF orthologs retain the capacity to recognize and bind vertebrate chromatin.

To better understand the chromatin binding dynamics of CTCF orthologs, we performed a clustering of the ChIP-seq signal across all experiments (**Figure 2C**). This analysis separated the CTCF ChIP-seq signal across four distinct clusters. Cluster 1, the most abundant one (56.70% of peaks), included chromatin regions only bound by the vertebrate CTCF orthologs. Cluster 2, also quite abundant (39.80%), involved binding events that are exclusive of chordate CTCF orthologs, that is for sea squirt, zebrafish and mouse. Cluster 3 represented a subset of chordate-specific events where the sea squirt CTCF ortholog displayed stronger binding capacities than its vertebrate counterparts (2.25%). Finally, cluster 4 denotes a minor subset of peaks (1,25%) that are bilateria-specific, with prominent binding for all orthologous versions except sea urchin, which binds to a lesser degree. A phylogenomic analysis on the CTCF ZFs recapitulated the bilaterian phylogeny. Yet, we observed that sea urchin ortholog has evolved rapidly in comparison to other bilaterian groups, which may explain its reduced capacity to bind to vertebrate genomes (**Supplementary Figure 6**). Insulation-score analysis of CTCF-associated clusters showed that invertebrate CTCF orthologs exhibit substantially weaker insulation than vertebrate CTCFs (**Supplementary Figure 7**). In fact, the insulation scores of invertebrate CTCF orthologs were comparable to those observed in the CTCF degron line treated with IAA (**Supplementary Figure 5**). This effect was particularly striking for the sea squirt CTCF ortholog which, despite binding pervasively across the vertebrate genome, was insufficient to establish robust chromatin insulation (**Supplementary Figure 7**). Collectively, these results demonstrate that while invertebrate orthologs can bind extensively across the vertebrate genome, chromatin binding per se is not sufficient to enable TAD formation via loop extrusion.

### CTCF has shifted its chromatin binding preferences during bilaterian evolution

To explore the sequence preferences of CTCF binding, we performed *de novo* motif discovery analysis independently for each ChIP-seq cluster (**Figure 2D**). For all clusters, the highest scored motif was similar to those previously described for CTCF, yet with some notable differences. Cluster 1, prominently bound by vertebrate CTCFs, and cluster 2, which was also bound by the sea squirt ortholog, were enriched for a motif reminiscent of that reported in vertebrates^46^. The motif associated with cluster 3, which was strongly bound by sea squirt CTCF, was also similar to the described vertebrate motif but lacking an adenine in a central position. Importantly, this nucleotide was highly scored in all other clusters or in motifs previously described for CTCF, denoting species-specific preferences. Cluster 4, bound by all studied species, was very similar to the motif previously reported for *Drosophila*^47^, lacking the 5’ domain that is characteristically bound by vertebrate orthologs. Strikingly, al clusters except the vertebrate-specific (cluster 1) were characterized by the presence of a 3′’terminal cytosine, indicating that this nucleotide may be critical for the binding of the invertebrate ortholog. As CTCF binding has been shown to be sensitive to DNA methylation (5mC), we hypothesized that the presence of this terminal cytosine may add an additional methylation site within the motif, thereby potentially reducing CTCF binding capacity. To explore this, we identified all potential instances of the invertebrate motif across the mouse genome and clustered those regions according to CTCF binding signal (**Supplementary Figure 8**). This analysis revealed clusters with distinct binding profiles. Cluster 1 and 2, which included bilateria and chordate-specific CTCF binding, tended to be evolutionary conserved and displayed minimal 5mC methylation. Cluster 3, which was the most abundant, comprised not conserved regions where CTCF was not bound. Interestingly, a subset of these regions was methylated at cytosines present in the motif, with the terminal one showing the highest 5mC levels (**Supplementary Figure 8**). These analyses suggest that the terminal cytosine associated with invertebrate CTCF is susceptible to undergo 5mC methylation in a vertebrate background, and that this pattern is associated with the absence of productive binding.

We next explored the genomic distribution of CTCF binding events across ortholog clusters and associated them with specific genomic features (**Figure 2E**). In clusters 1 and 2, corresponding to vertebrate-specific and chordate-specific CTCF-bound regions, most binding events were located in distal regions (gene bodies or intergenic regions), with a smaller proportion occurring proximal to promoters. In contrast, the proportion of promoter-proximal binding events increased notably in clusters associated with invertebrate CTCF orthologs, namely the sea squirt- and bilaterian-specific clusters (3 and 4, respectively). These results suggest that CTCF has a preferential bias towards distal genomic regions in vertebrates. Yet, this distal bias may have emerged earlier in the chordate lineage, as it was already apparent in cluster 2, which includes binding events for the sea squirt ortholog. Collectively, our results indicate that CTCF acquired distinct DNA-binding motifs and genomic distribution during bilaterian evolution.

### The vertebrate CTCF protein is not sufficient to form TADs via loop extrusion

Having established that a vertebrate version of CTCF is required for TAD formation through loop extrusion in mESCs, we next asked whether it is also sufficient to mediate this process. We explored this aspect by replacing the endogenous invertebrate CTCF by its vertebrate mouse ortholog, in a fly background. In *Drosophila*, loop extrusion is mainly involved in enhancer promoter interactions, with a limited contribution to chromatin domain formation^23^. Furthermore, CTCF is dispensable for embryonic development in *Drosophila*, in contrast to mouse^42^. Therefore, we attempted to generate viable flies carrying homozygous CTCF replacements. For this purpose, we engineered a fly line (**Figure 3A**) carrying an N-terminal FLAG-Tag, followed by an FRT recombinase site, and the native *D. mel. CTCF* gene. A GFP marker indicates the presence of the transgenic construct. Another FRT site, oriented as the first, was located downstream of the GFP, followed by the mouse *Ctcf* CDS and the fly 3′UTR. Upon crossing with a constitutive *FLP* line, the endogenous *D. mel*. *CTCF* gene and the GFP marker are excised via FRT recombination, leaving the mouse ortholog in frame with the FLAG-tag and expressed under the regulation of the fly promoter, 5′UTR and 3′UTR (**Figure 3B**). Western blot analysis using an anti-FLAG antibody confirmed the expression of the species-specific CTCF protein, with their corresponding expected sizes (**Figure 3B**). Homozygous replacement fly lines expressing the mouse CTCF version were viable, as also described for CTCF KO (*CTCF0*)^42^. Mutant flies displayed a reduced lifespan and did not survive beyond day 4 after eclosure. Flies expressing the mouse CTCF displayed the *Hox* phenotypes characteristic of the CTCF KO, as well as abnormal wing position and reduced ability to fly (**Figure 3C**). Protruding genitalia and abnormal pigmentation were also observed in males (**Figure 3C**). Further, mutant animals displayed neurological phenotypes, as also described for CTCF KO mutants^42^, including spasmatic and uncoordinated movements. Locomotion climbing assays performed 8–10 hours after eclosion revealed that flies carrying the mouse ortholog exhibited significantly reduced speed and gravitaxis compared with controls carrying the fly CTCF (**Supplementary Figure 9**). Moreover, mouse CTCF flies rapidly lost their ability to walk or climb as they aged. These results suggest that the replacement of *Drosophila* CTCF by its mouse ortholog counterpart mainly results in loss-of-function phenotypes in mutant flies.

**Fig. 3.**
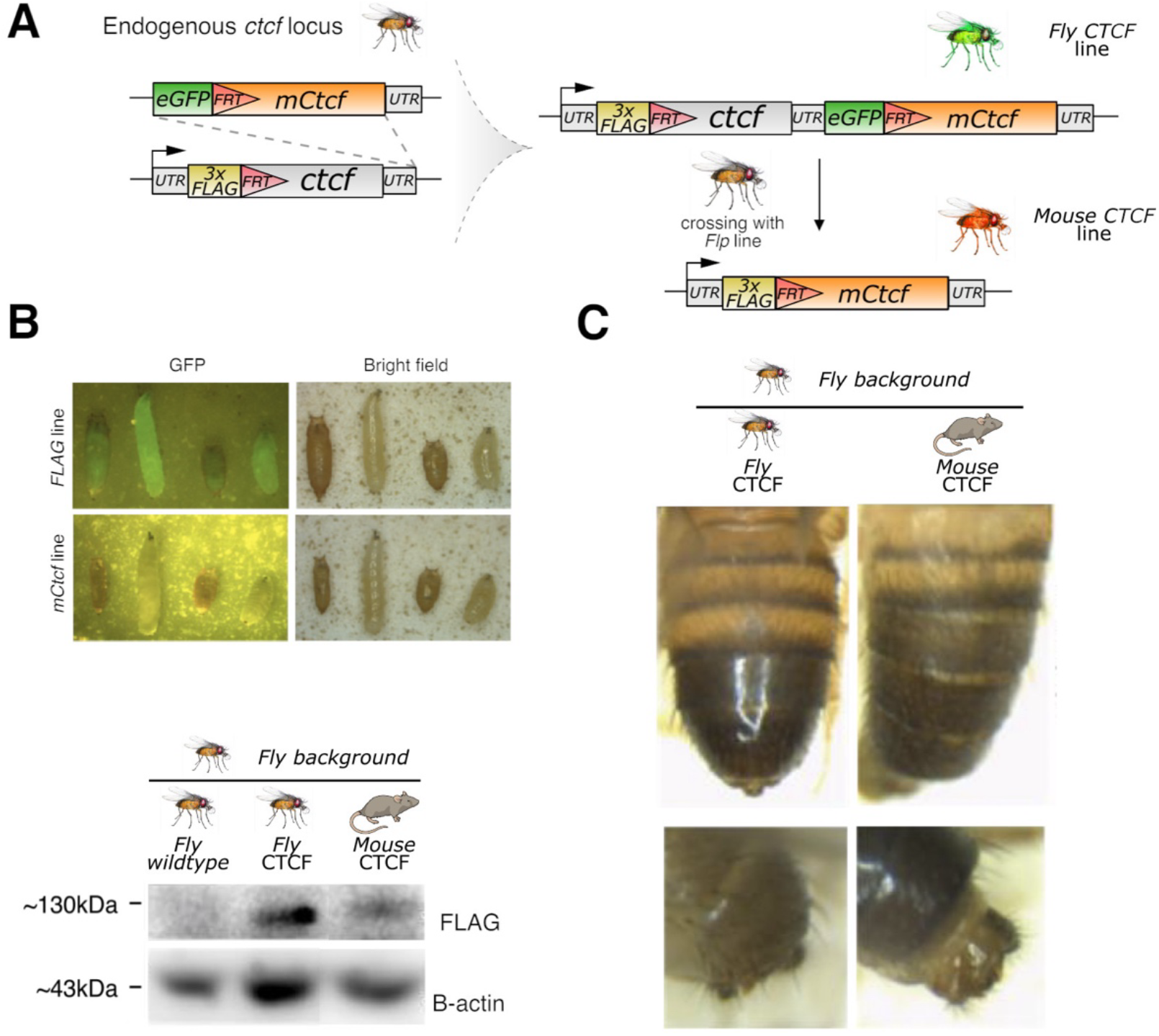
CTCF orthologous replacement in flies. **A.** Experimental strategy for generating flies carrying an orthologous replacement of CTCF. First, fly embryos were engineered to introduce a FLAG-tag followed by an FRT site at the endogenous CTCF locus. Subsequently, these embryos were modified to insert a construct containing an GFP marker, a second FRT site, the mouse Ctcf coding sequence (CDS), and the fly 3′UTR downstream of the endogenous CTCF gene. Upon crossing to a fly line expressing FLP recombinase, the endogenous fly CTCF coding sequence was excised through FLP/FRT-mediated recombination, leaving the mouse CTCF ortholog, fused to the N-terminal Flag tag, under the control of the endogenous fly ctcf promoter and 3′UTR. **B.** Top, fruit fly larvae before and after FLP-mediated recombination. The difference in body size reflects the presence or absence of the Tb (Tubby)-marked balancer chromosome. Larvae carrying the balancer (heterozygous for the CTCF construct) display the characteristic small-bodied Tubby phenotype, whereas larvae lacking the balancer (homozygous for the CTCF construct) exhibit a normal body size. Bottom, Western blot analyses on mutant controls and mouse replacement flies using a common anti-FLAG antibody. Note the presence of a specific band with the expected sizes for both replacement lines. The image has been modified to remove an unrelated lane (see Methods) **C**. Abdominal and genital phenotypes in mutant male flies.

Next we evaluated the effects of CTCF orthologous replacement on 3D chromatin organization. We performed Hi-C experiments on the central nervous system (CNS) from the third instar larval stage, a tissue that has been extensively characterized in this context^43^ (**Figure 4A**). Eigenvector analyses revealed that A/B compartmentalization was largely unaffected in flies carrying a mouse CTCF replacement (**Supplementary Figure 10**). We subsequently determined chromatin domains by calculating insulation scores. This analysis showed that most boundaries retained their insulation capacity (**Figure 4B** and **Supplementary Figure 11**). Yet, a small subset of boundaries lost insulation in mouse CTCF-replacement mutants compared to fly control embryos (**Figure 4C**). Importantly, those affected boundaries corresponded to the same regions that were previously identified as altered in CTCF KO mutants^43^ (**Supplementary Figure 12**), confirming again that the effects associated with introducing the mouse ortholog are largely derived from loss-of-function effects. Crucially, we could not detect any region that gained insulation in mutants, or any *de novo* formation of chromatin loops. These results indicate that the vertebrate CTCF protein is not sufficient to form TADs via loop extrusion in an invertebrate background. Moreover, the vertebrate CTCF orthologous also cannot perform the functions associated to its invertebrate counterpart, which acts as an insulator protein on certain boundary regions or regulator of *Hox* genes^42,43^. Collectively, these findings demonstrate that the invertebrate and vertebrate CTCF orthologs have followed divergent functional paths during evolution.

**Fig. 4.**
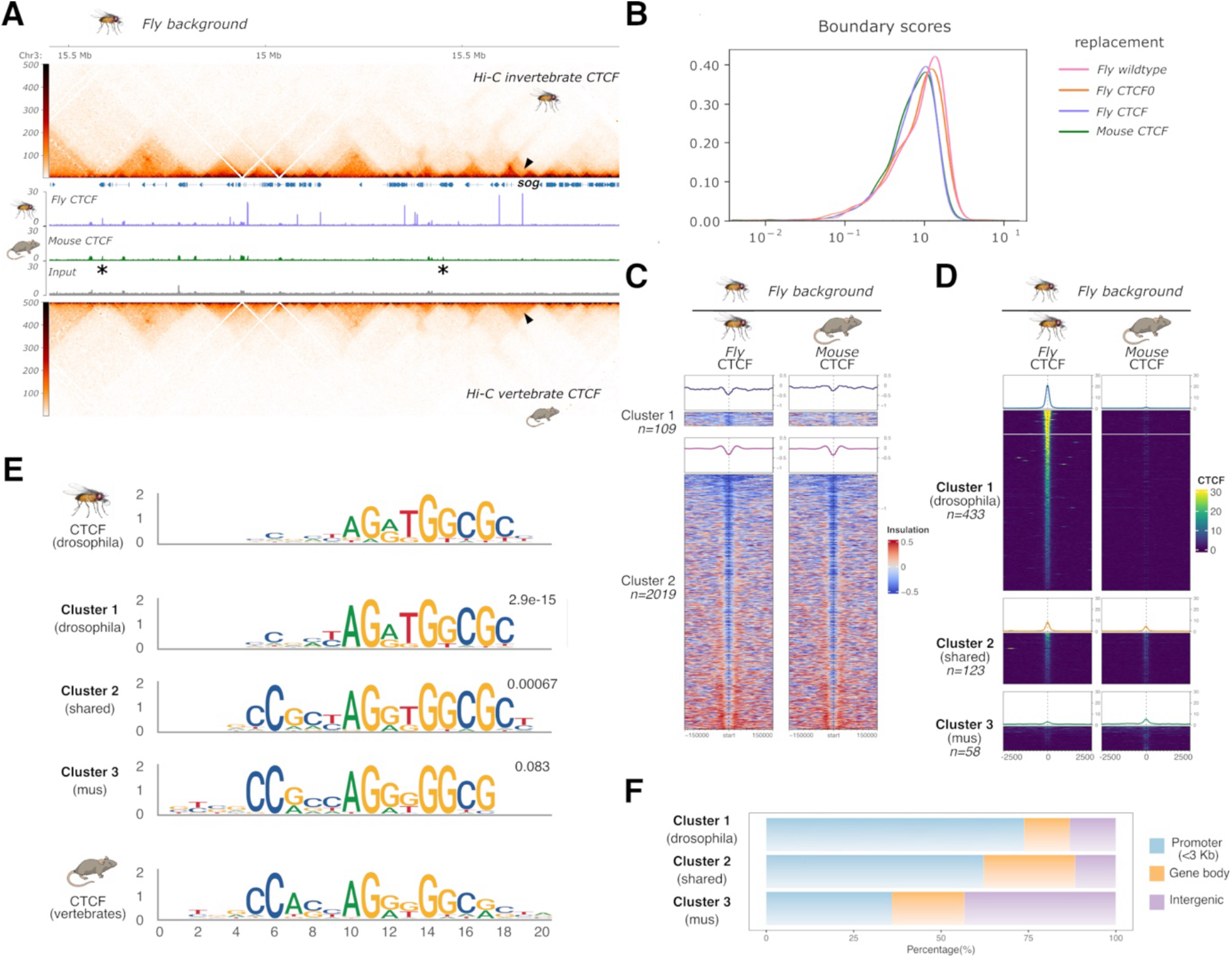
Functional effects of CTCF orthologous replacement in Drosophila mutants. **A.** Effects of CTCF replacement in the vertebrate chromatin in a representative genomic region. Top and bottom, Hi-C maps from CNS of fly control and mouse orthologous replacements, respectively. Black arrow denotes a differential TAD boundary among both samples. Middle, ChIP-seq profiles of CTCF ortholog binding in CNS from mutant lines. Note the absence of new chromatin domains or extrusion trails, despite the binding of mouse CTCF (asterisks). All ChIP-seq experiments were performed using a common antiFLAG antibody, making them directly comparable. **B.** Distribution of boundary scores from Hi-C experiments. Note that all samples show a similar distribution. **C.** Clustering of genomic regions according to insulation signal **D.** Clustering of CTCF ChIP-seq signal at specific genomic regions. **E.** Most represented motif per ChIP-seq cluster. The canonical CTCF motif described for vertebrates and Drosophila is included for reference. **F.** Genomic distribution of CTCF binding events per each ChIP-seq cluster.

### Vertebrate CTCF exhibits a distal binding bias across the invertebrate genome

Given the limited effects on chromatin organization associated with the mouse CTCF protein when introduced in flies, we explored if this ortholog can effectively bind across the invertebrate genome. We leveraged the introduced FLAG-tags to perform ChIP-seq experiments in CNS, using a common antibody for both mouse CTCF replacement mutants and fly control lines (**Figure 4A**). The inspection of ChIP-seq profiles denoted widespread binding events for both orthologs at the genome-wide level, confirming that the vertebrate CTCF can also bind on invertebrate genomes. We subsequently performed clustering analyses using the combined ChIP-seq signal from both experiments (**Figure 4D**). This analysis revealed that most binding events correspond to regions associated with the *Drosophila* CTCF, but not with its mouse counterpart (Cluster 1, 70.52%). A considerable number of genomic regions where also bound by both orthologous proteins (Cluster 2, 20.03%). Interestingly, we also find chromatin locations were the mouse ortholog appears to bind with more affinity than the fly counterpart (Cluster 3, 9.45%). Insulation scores revealed a decrease in signal on Cluster 1 regions for mouse CTCF replacement flies, consistent with the observed lack of binding of this ortholog (**Supplementary Figure 13**). Cluster 2, which features binding sites for the two CTCF orthologs, did not show noticeable changes in insulation between conditions (**Supplementary Figure 13**). This suggests that the reduced insulation observed in some boundaries of the mouse CTCF replacement line may derive from the lack of binding rather than protein effects. For Cluster 3, which includes regions that are more prominently bound by the mouse protein, we do not observe any changes in insulation, neither the formation of new boundaries (**Supplementary Figure 13**). These results again confirm that the effects of the mouse CTCF protein on 3D chromatin organization are relatively minor in the invertebrate genome.

Finally, we explored the binding preferences of CTCF orthologs in the invertebrate genome. *De novo* motif discovery revealed that Cluster 1 was enriched for a short motif analogous to the one previously described in *Drosophila*^47^ (**Figure 4E**). Cluster 2, which is bound by both orthologs, was enriched for the extended motif that is similar to the one reported for vertebrates^46^ (**Figure 4E**). However, this motif featured the additional 3′ terminal cytosine that is characteristic of invertebrate CTCF, as also identified in our mESC replacement experiments (**Figure 2C**). The motif identified in Cluster 3, which comprises genomic regions more strongly bound by mouse CTCF, also contained the extended vertebrate-like motif. Yet, it lacked the 3′ terminal cytosine, further supporting its importance for invertebrate CTCF binding (**Figure 4E**). The genomic distribution of binding events across clusters also mirrored the patterns observed in the mESC replacement experiments (**Figure 4F**). Clusters preferentially bound by the invertebrate CTCF protein were enriched at promoter-proximal regions, whereas the vertebrate-specific cluster was predominantly associated with distal genomic regions. Together, these findings are consistent with our previous observations and support the evolutionary diversification of both the CTCF binding motif and its genomic binding preferences across bilaterian evolution.

### CTCF functions in transcriptional regulation and chromatin folding are independent and evolutionary uncoupled

Despite the notable differences in architectural function observed between vertebrate and invertebrate CTCF versions, some orthologs were able to bind extensively across their non-native genomes (**Figure 2A, 2B, 4A and 4B**). Given the multifaceted nature of CTCF, we sought to explore whether the presence of orthologous CTCF proteins may affect the transcriptional landscape, independently of 3D chromatin organization. We performed RNA-seq experiments, focusing on our replacement mESC lines, given the extensive binding of sea squirt CTCF across the vertebrate genome and its inability to form TADs via loop extrusion. Upon addition of IAA, we observe 534 dysregulated genes in the control line, consistent with previous reports^15^ (**Supplementary Figure 14, 15 and 16**). Importantly, we observe that a percentage of those genes were rescued upon the expression of invertebrate CTCF orthologs. Notably, the highest number of rescued genes (124 in total) was observed upon the expression of the sea squirt CTCF. Focusing on genes associated with promoter CTCF binding, we observed that a large number of those displaying vertebrate-specific binding were effectively rescued (clusters 1 and 2 in **Supplementary Figure 14**). This rescue effect was more pronounced for downregulated than upregulated genes, suggesting that CTCF may predominantly act as a transcriptional activator in mESC when bound near promoters. Interestingly, we observed a similar effect for the sea squirt CTCF, which again showed the highest degree of rescue among the invertebrate orthologs, and specifically for downregulated genes associated proximal CTCF binding sites (cluster 2 in **Supplementary Figure 14**). One notable example of such rescue is *Ints1*, a gene encoding a subunit of the Integrator complex and whose depletion has been previously shown to cause cell death in mESCs^48,49^. Specifically, the expression of chordate orthologs, that is either sea squirt, zebrafish, or mouse, led to CTCF binding at the *Ints1* promoter and restored expression to normal levels (**Supplementary Figure 14 and 16**).

These observations prompted us to explore if the introduction of the sea squirt ortholog may compensate for the lethality associated with the lack of endogenous CTCF in mESC^41^. To address such question, we employed a similar CRISPR-mediated homologous recombination strategy but aimed to generate homozygous replacement cell lines directly (**Figure 5A**). The replacement of mouse CTCF with the *Drosophila* ortholog yielded no viable homozygous mESC clones, suggesting that such replacement leads to loss-of-function effects and cell lethality. In contrast, the CRISPR-mediated replacement with the Ciona ortholog yielded homozygous knock-in mESC clones at a frequency of >40% (**Figure 5A**). Despite lacking endogenous mouse CTCF expression (**Figure 5B**), homozygous clones could be maintained in culture and serially passaged, although they proliferated slower than control cells (**Figure 5C**). These findings demonstrate that the expression of the sea squirt ortholog is sufficient to rescue the lethality associated with the loss of the endogenous CTCF in mESC.

**Fig. 5.**
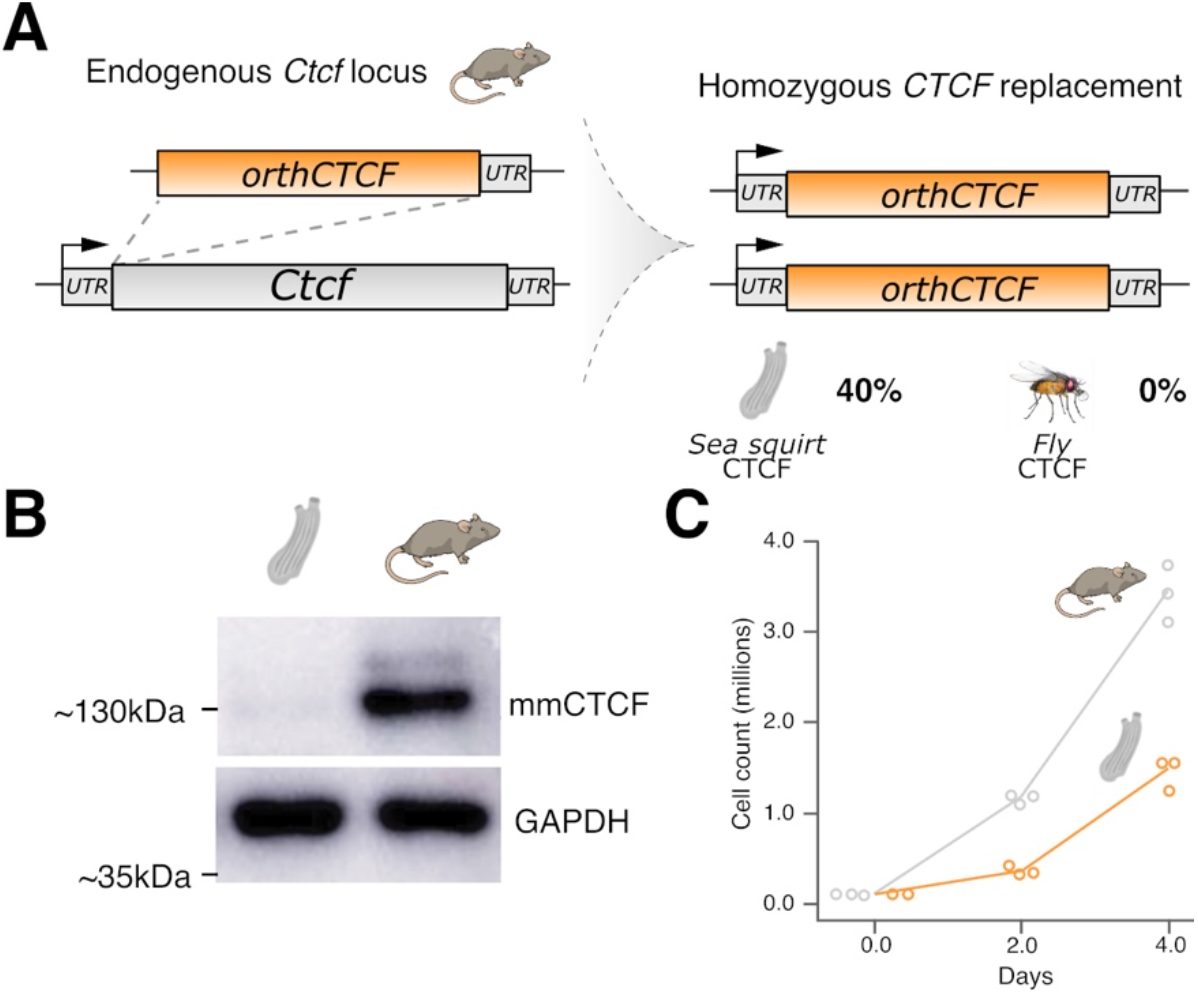
Expression of sea squirt ortholog rescues viability in CTCF knockout mESC cells. **a.** Experimental strategy to generate mutant mESC carrying homozygous CTCF orthologous replacements. Direct replacements were performed using sea squirt CTCF yielded more than 40% mESC clones with homozygous replacements, while no equivalent clones were obtained when using the fly ortholog. **b.** Western blot analyses on mESC carrying homozygous sea squirt replacements showing the absence of the endogenous CTCF protein. **c**. Growth curves for mESC carrying homozygous sea squirt replacements and mouse controls. While sea squirt replacement mESC grow slower than mouse controls, they can be effectively cultured and passaged.

Altogether, our findings support the existence of two genetically separable functions of CTCF, one as a transcriptional regulator and another as a mediator of 3D genome organization through loop extrusion. Remarkably, these functions also appear to be evolutionarily uncoupled, with the transcriptional role of CTCF showing conservation across chordates, and the architectural function apparently limited to vertebrate species.

### The function of CTCF as a loop extrusion barrier was present in the last common ancestor of all vertebrates

Having established that the distinct functions of CTCF are uncoupled, we next sought to determine when its capacity to act as a barrier to loop extrusion emerged during evolution. In a previous study, we could show that signatures of loop extrusion are already observable in the genome of the little skate *Leucoraja erinacea*^9^, placing the emergence of CTCF-mediated loop formation, at the latest, at the origin of gnathostomes (jawed vertebrates). To further refine the evolutionary timing of this event, we performed additional chromatin conformation analyses in cyclostomes, a key lineage to understand the origin of vertebrates (**Supplementary Figure 17**). Hi-C experiments in adult liver tissue from the sea lamprey, *Petromyzon marinus,* revealed that its genome exhibits a type II architecture, with chromosomes organized into distinct territories within the nucleus (**Supplementary Figure 18**). At higher resolution, lamprey chromosomes were segregated into distinct compartments, with the A compartment displaying higher gene density than the B compartment (**Supplementary Figure 19**). At the sub-megabase scale, the lamprey genome was organized into TADs (**Figure 6A)**, with their boundaries showing evidence for clusters of CTCF binding sites with a divergent motif orientation (**Figure 6B**).

**Fig. 6.**
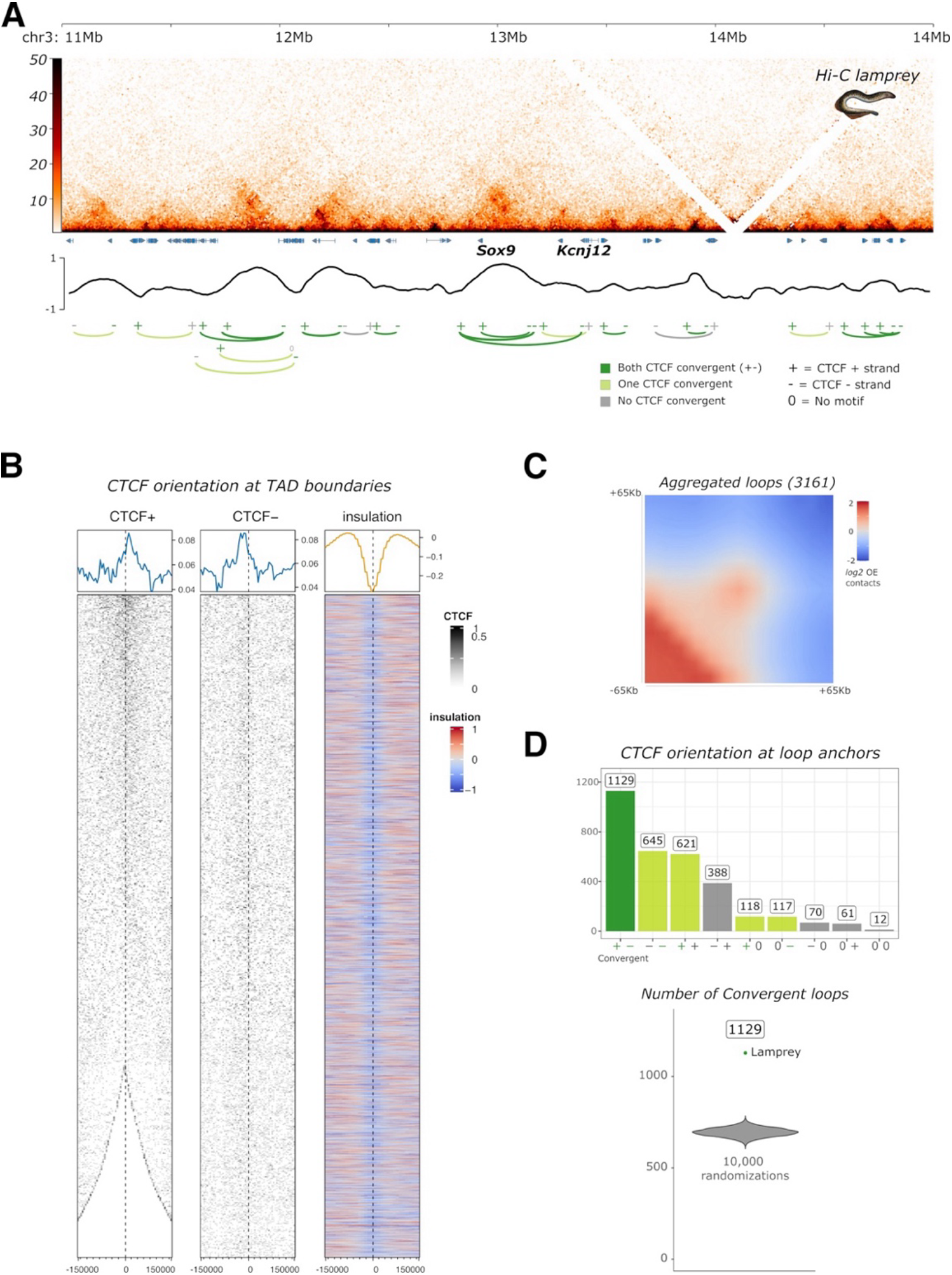
3D chromatin organization in cyclostomes. **a.** KR normalized Hi-C heatmap of sea lamprey (Petromyzon marinus) liver samples, showing a representative locus at 5 kb resolution. Genes and insulation scores are indicated below. Chromatin loops are indicated with arches that are color coded according to the orientation of the best CTCF motifs found at the anchors (dark green: both CTCF convergent, light green: one CTCF convergent, grey: no CTCF convergent). The orientation of the best CTCF motif at the corresponding anchors are displayed. **b**. Clustering of CTCF motif orientation at TAD boundaries. **c.** Aggregated observed/expected (OE) contacts around the 3161 chromatin loops identified in lampreys. **d.** Orientation of chromatin loops. Top, barplot quantifying the abundance of chromatin loops classified by the orientation of the best CTCF motif at their anchors showing. Fully convergent loops are most abundant (randomized p-value < 10^-5^, see Methods). Bottom, randomization analysis showing the expected number of convergent loops under a random model. The observed number exceeds that expected by chance. A total of 10,000 randomizations were performed.

Genome-wide analysis of chromatin loops revealed that this feature is pervasively found across the lamprey genome (**Figure 6A** and **C**). Motif analyses showed that nearly half of the detected loops were associated with CTCF motifs displaying a convergent orientation (**Figure 6C**). An additional fraction of loops contained one anchor associated with a CTCF motif oriented towards the opposing anchor, whereas the corresponding motif at the second anchor was not detectable. Some instances of CTCF motif convergency can be observed at developmental loci such as the *Sox9* TAD (**Figure 6A**), where the orientation and location of boundary and promoter-anchored loops is similar to those observed in mammals (**Figure 2A**). Collectively, these results suggest that 3D chromatin organization in lampreys arises from the interplay of two mechanisms, compartmentalization and TAD formation mediated by loop extrusion, resembling the genome organization described in jawed vertebrates. Furthermore, the predominant presence of loops with motifs in a convergent orientation suggests that the ability of CTCF to modulate loop extrusion was already present in the common ancestor of all vertebrates.

## DISCUSSION

Here we explored how the functions of CTCF have diversified during evolution and the potential implications at the regulatory level. Through replacement experiments in mouse and fly, we found evidence of a substantial functional divergence between invertebrate and vertebrate orthologs, particularly in the ability of CTCF to shape 3D genome organization. In *Drosophila*, CTCF contributes to 3D genome organization, as its depletion results in the loss of a subset of domain boundaries^43^. However, this function operates within a broader and partially redundant network of insulator proteins^50–53^, suggesting that CTCF is just another component of the architectural machinery rather than a central organizer. This is consistent with the relatively limited effects of cohesin perturbation on global domain organization in flies, despite some noticeable effects on enhancer-promoter interactions^23^. In contrast, CTCF has a much more prominent role in vertebrate 3D genome organization, highlighted by a global disappearance of TADs and loops that is similar to the effects associated with the disruption of loop extrusion^15,16^. Our experimental setup provides evidence that the vertebrate version of CTCF is necessary to form TADs via loop extrusion. Yet this factor alone is not sufficient to induce their formation of such domain in an invertebrate background, indicating the existence of additional requirements to support this function. Consistent with recent studies^54^, our results also provide a clear separation between CTCF functions, denoted by the ability of sea squirt CTCF to rescue the viability of mESCs in the absence of TAD formation. This observation may also imply that the role of CTCF to act as a transcriptional regulator is evolutionarily ancient relative to its specialized role as a barrier for loop extrusion. In agreement with this separation of functions, recent work in organoid models has shown that CTCF-dependent looping is dispensable during early developmental stages but becomes essential at later stages, indicating differential roles during development^55^. Therefore, while CTCF may already have certain capacity to contribute to 3D genome organization in early metazoans, it subsequently acquired a more central role during evolution through a tight integration with the loop extrusion machinery.

The functional diversification of CTCF seems to be linked to the evolution of its N-terminal domain, which plays a fundamental role in its interaction with cohesin^37,38^. A critical sequence within this region is the YXF motif, which is required for CTCF–cohesin interaction and is conserved across bilaterians^37,38^. This motif is present in species such as fly, sea urchin or sea squirt, where CTCF does not seem to function as a prominent barrier to cohesin-mediated loop extrusion. This raises the possibility that the ability of CTCF to interact with cohesin may be an ancient feature that predates the emergence of its architectural role in vertebrates. In fact, the interplay between CTCF and cohesin in genome organization extends beyond interphase^56,57^. An additional N-terminal motif, KTYQR, shows a much more restricted distribution across bilaterian lineages, being conserved primarily among jawed vertebrates^43^. Recent *in vitro* experiments have shown that the KTYQR, together with the YXF motif, contributes to the ability of the CTCF N-terminal region to impede cohesin-mediated loop extrusion, with the two motifs acting through distinct but complementary mechanisms^36^. Interestingly, the KTYQR motif is absent from the sea lamprey CTCF protein, which has undergone a prominent divergence at its N-terminal domain. The fact that signatures of loop extrusion are readily observed in this species suggests that the molecular mechanisms underlying CTCF-mediated looping may have evolved through the modification of multiple sequence features, rather than through the acquisition of a single motif. This finding is consistent with a recent study employing a plasmid-based approach, which demonstrated that the CTCF from the Siberian brook lamprey (*Lethenteron kessleri*) can support loop formation in human HCT116 cells^58^. Therefore, the YXF and KTYQR motifs may not account for all functions of the CTCF N-terminal domain, suggesting that additional sequence features yet to discover may also contribute to cohesin recruitment, retention, or blocking.

Our data support a model in which the evolution of CTCF interactions with cohesin or with DNA may have followed distinct trajectories. Recent research suggests that N-terminal domain diversification underlies a gradual increase in the ability of CTCF to support loop formation across bilaterian evolution^58^. Here we identify a substantial shift in DNA-binding preferences near the base of the chordate lineage. When introduced into a vertebrate background, the sea squirt CTCF binds extensively across the genome, sharing a substantial number of sites that are also recognized by vertebrate CTCF. Importantly, these sites are associated with the expanded motif that is characteristic of vertebrates^46^, featuring two cytosines at the 5′end. However, despite this pervasive binding and motif similarities, the sea squirt protein also shows additional signatures of sequence preferences that are observed in invertebrate CTCF orthologs. In particular, its binding appears to depend on the presence of a 3′terminal cytosine, a feature also observed for sea urchin or fly CTCF, the last one irrespective of the genomic background in which it is expressed. The conservation of this 3′terminal cytosine is particularly intriguing, as it generates an additional site that is susceptible to 5mC methylation, as observed in mESC. Since 5mC is a more prevalent mechanism in vertebrates^59^ and can impair CTCF binding^60–62^, evolution may have selected against the dependence on this terminal cytosine in this clade, favoring a diversification of the CTCF binding motif.

Beyond nucleotide preferences, we also observe a remarkable transition in the genomic distribution of CTCF binding sites, with a preference for distal sites in vertebrates in comparison to invertebrates. Interestingly, some studies evidence that promoter-proximal and distal binding sites may constitute functionally distinct classes^54,63,64^, indicating that the “CTCF code” is probably more complex than previously anticipated. It is tempting to speculate that the distalization of CTCF binding observed in the vertebrate clade may be related to the evolution of its DNA-binding domain. In a mouse background, we observed that invertebrate CTCF orthologs bind preferentially to promoter regions, which are among the few regions that remain largely unmethylated in vertebrates^65^. One possibility is that, in the vertebrate context, the methylation of the 3′ terminal cytosine within the CTCF motif may limit the binding of invertebrate CTCF proteins to distal regions, thereby favoring their occupancy at promoter-proximal sites. Consistent with this hypothesis, the sites that are more strongly bound by the sea squirt CTCF on the vertebrate genome correspond predominantly to promoter-associated sites. Yet, those still represent only a small fraction of all sites that can be recognized by the protein, as most of the binding events are located in distal genomic regions. This indicates that sea squirt CTCF has an intrinsic capacity to recognize distal elements, suggesting that the distalization of CTCF binding already occurred during early chordate evolution. Importantly, this preference for distal binding would have been difficult to infer from studies directly on the sea squirt, given its compact genome and the limited availability of distal regulatory space in tunicates^66^. In this respect, further studies of other early-branching chordates with non-compacted genomes may shed light on the molecular basis of this distalized binding pattern. Nevertheless, exchanging orthologous into controlled genomic backgrounds provide valuable means to disentangle the intrinsic properties of a protein from the evolutionary constraints imposed by their native contexts.

Our chromatin studies in cyclostomes provide evidence that the capacity of CTCF to act as a barrier to loop extrusion was already present in the last common ancestor of all vertebrates, at least after the first round (1R) of Whole Genome Duplication shared by cyclostomes and jawed vertebrates^67^ (**Supplementary Figure 17**). A fundamental question therefore arises: which regulatory functions may have been enabled by this ancestral architectural mechanism? One possibility is that CTCF-mediated looping allowed to cope with the increasingly complex regulatory landscapes that emerged during vertebrate evolution. Vertebrate genomes contain extensive repertoires of tissue- and developmental-stage-specific regulatory elements^68^, with regulatory interactions frequently occurring over distances of tens to hundreds of kilobases. Accumulating evidence suggests that, in vertebrates, cohesin-mediated loop extrusion is particularly important for regulatory interactions over longer genomic distances (>20–50 kb), whereas many proximal enhancer–promoter interactions persist following the disruption of this mechanism^69–74^. In such context, CTCF allows the rewiring of loop extrusion trajectories, providing means for the stabilization of distal regulatory interactions^75^. However, a detrimental effect of such regulatory expansion would be the increased potential for aberrant interactions between regulatory elements and genes. In such context, the clustering of CTCF sites at domain boundaries provides a complementary mechanism to partition an expanded regulatory space, constraining the genomic range over which regulatory elements can interact. Thus, the emergence of a CTCF–mediated looping system may have provided both an opportunity and a constraint: it could promote distal regulation while simultaneously imposing architectural boundaries that restrict inappropriate contacts. This model highlights an important contrast with invertebrates species such as *Drosophila*, where extreme cases of long-range regulatory interactions above the megabase scale have been reported^76^. Therefore, extreme regulatory distances are not intrinsically incompatible with invertebrate genome organization. Rather, our findings suggest that vertebrate evolution may have adopted a distinct architectural solution for managing an increasingly complex regulatory landscape. Importantly, we find evidence that the distinct ancestral molecular properties of CTCF, such as DNA binding or cohesin interaction, may have been progressively modified into this elaborated architectural system. Thus, rather than inventing completely new functions, evolution may have reshaped ancient molecular capabilities, enabling CTCF to become a central organizer of the 3D genome.

## ACKNOWLEDGEMENTS

We thank the Genomics Technology Platform of the Max Delbrück Centre for Molecular Medicine, particularly Madlen Sohn and Tatiana Borodina, for technical assistance. We thank Eileen Furlong, Maria Cristina Gambetta and Robert Hiesinger for sharing fly stocks. We thank members of the Lupiáñez laboratory for fruitful discussions. This project was funded by Deutsche Forschungsgemeinschaft via the Priority Programs 2202 (project no. 422856854; LU 2426/2-1 to D.G.L, and ZI 1517/5-1 to R.Z). Research in the Lupiáñez lab was funded by the European Research Council (grant no. 101045439, 3D-REVOLUTION) and by the Spanish “Agencia Estatal de Investigación” (grant number PID2022-143253NB-I00/ AE,I/10.13039/501100011033/ FEDER, UE). Funded by the European Union. Views and opinions expressed are however those of the author(s) only and do not necessarily reflect those of the European Union or the European Research Council Executive Agency. Neither the European Union nor the granting authority can be held responsible for them. Work in the L.C. lab at NYU was supported by NIH awards R01 HL108643 and R01 HD096770. L.A. was supported by a fellowship from the MDC-NYU PhD Exchange Program. R.D.A. was supported by an EMBO Postdoctoral Fellowship (grant no. EMBO ALTF 537-2020). AMR acknowledges support from PID2024-162736OB-I00 and to European Commission’s Horizon Europe Research and Innovation program (OSCARS, No. 101129751).

## AUTHOR CONTRIBUTIONS STATEMENT

### Contributions

L.A., R.Z, and D.G.L. originally conceived the study and designed the experiments, with subsequent contributions from R.D.A. L.A. designed mESC and fly replacement strategies, with input from R.Z and D.G.L. L.A. and H.S generated, validated mESC replacement lines and collected material. L.A. performed Hi-C, ChIP-seq and RNA-seq experiments on mESC lines. L.A and R.D.A. analyzed mESC omics data. H.S. performed fly husbandry, molecular and phenotypical characterization, as well as RNA-seq experiments. L.A. performed Hi-C and ChIP-seq experiments in fly replacement lines. L.A and R.D.A. analyzed fly omics data. C.R. and L.A performed experiments in sea squirt, with input from L.C. L.A and R.D.A. analyzed sea squirt omics data. J.R, O.B. and F.M.R. performed experiments in sea lamprey. R.D.A. analyzed sea lamprey omics data. AMR conducted sequence identification and phylogenetics analyses. R.D.A. and D.G.L. wrote the manuscript with input from all authors.

## MATERIAL AND METHODS

Research carried out in this study complies with all relevant ethical regulations for animal experimentations. Sea lampreys (Petromyzon marinus) captured in the River Miño by professional fishermen were obtained from PESBAMI, S.L. (Torreiro 15, Soutelo, Pontevedra, Spain), a company authorized to commercialize this species. Animals were transported to the facilities of the Instituto de Investigaciones Marinas (IIM-CSIC), where they were euthanized by immersion in an overdose of MS-222 (500 mg L⁻¹ for 20–30 min; Sigma-Aldrich, USA) before biological samples were collected. All procedures were conducted under the ethical authorization ES360570202001/18/FUN.01/BIOL.AN.08/JRM, granted by the Institutional Animal Care and Use Committee of the IIM-CSIC, in accordance with Spanish legislation on the protection of animals used for scientific purposes (Royal Decree 53/2013) and European Directive 2010/63/EU.

### Generation of mESC carrying CTCF orthologous replacements

mESCs were maintained in ESC medium supplemented with leukemia inhibitory factor (LIF; 1:10,000) and cultured in 6-cm dishes on a layer of mouse embryonic fibroblasts (MEFs). ESC medium was prepared by supplementing 625 mL of basal medium with 93.75 mL fetal calf serum (FCS), 6.25 mL each of 100× non-essential amino acids, 100× glutamine (200 mM), 100× nucleosides, 10 mM β-mercaptoethanol, and penicillin (5,000 U/mL), together with streptomycin (5,000 U/mL). Cells were routinely passaged every 24–72 h by trypsinization and transferred to 6-cm dishes pre-coated with 0.1% gelatin. Cultures were maintained at 37°C, and the medium was replaced daily whenever passaging was not performed.

For genome editing, 5 × 10⁵ OsTir1/CTCF-AID mESCs were plated onto MEF feeder layers in 6-cm dishes. Cells were subsequently transfected with CRISPR constructs using the FuGENE transfection system (Promega), with 8 μg of sgRNA/nuclease construct and 4 μg of donor DNA. At 48 h post-transfection, the cells were passaged onto puromycin-resistant MEF feeder layers and subjected to puromycin selection for 48 h. Following selection, the surviving cells were allowed to recover and expand for 3–5 days before being transferred to 96-well plates containing MEF feeder cells. After 2–4 days of culture, each plate was replicated into three plates: two for cryopreservation and one for cell expansion and genomic DNA isolation. For cryopreservation, cells were washed twice with 1× PBS and detached by incubation with trypsin at 37°C for 5–7 min. Trypsin activity was subsequently neutralized by adding normal medium consisting of 80% KnockOut Dulbecco’s Modified Eagle Medium (KO-DMEM) and 20% FCS. The cell suspension was collected in a Falcon tube, and cell density was determined using a hemocytometer by counting a 10-μL aliquot. Cells were then pelleted by centrifugation at 1,000 × g for 5 min and resuspended in normal medium. Aliquots containing 5 × 10⁵ or 1 × 10⁶ cells were distributed into two or three cryogenic vials and mixed with freezing medium composed of 60% KO-DMEM, 20% FCS, and 20% dimethyl sulfoxide (DMSO). The vials were placed in CellHome freezing containers (Atlantis Bioscience) and subsequently stored at −80°C.

Genomic DNA from candidate mutant mESC clones was analyzed by PCR to verify the CRISPR/Cas9-mediated integration of the donor template. Amplification was performed using DreamTaq DNA polymerase (Thermo Fisher Scientific) and the primer pairs listed in **Supplementary Table 1**. Initial screening involved separate PCR reactions targeting the 5′ and 3′ junctions of the inserted sequence. Clones yielding positive amplification were subsequently analyzed using primers spanning the wild-type locus to establish the zygosity of the integration. The identity of the edited sequences was further confirmed by Sanger sequencing.

### IAA treatment of mESC

Cells were prepared for collection following a five-day experimental protocol. Heterozygous CTCF ortholog-replacement mESCs (5 × 10⁵ cells per replicate) were plated in duplicate on MEF feeder layers in 6-cm dishes and cultured for 48 h. To induce degradation of endogenous CTCF expressed from the wild-type allele, IAA was added to a final concentration of 500 μM for the final 24 h before harvesting. Cells were then detached by trypsinization, and residual MEFs were removed by sequentially allowing the cell suspensions to settle on gelatin-coated dishes for two 30-min periods. The resulting cell suspensions were collected and centrifuged at 1,000 rpm for 5 min. The cell pellets were retained for subsequent downstream analyses.

### Generation of mutant flies carrying CTCF orthologous replacements

Flies were kept at either 18°C or 25°C under a 12-h light cycle and at 60% relative humidity. Stocks were routinely transferred to fresh vials every 2–3 weeks and maintained on standard fly medium. The food was prepared in 14 L batches containing 192 g agar, 360 g dried yeast, 200 g soy powder, 440 g sugar syrup, 1,600 g malt extract, 1,600 g corn powder, 125 mL propionic acid, and 48 g nipagin.

Transgenic flies expressing the *Mus musculus* CTCF ortholog were generated using FLP/FRT-mediated site-specific recombination. The transgenic founder lines, generated by WellGenetics Inc. in a *w[1118]* genetic background, carried two engineered FRT sites at the endogenous *ctcf* locus. One FRT site was positioned downstream of a 3XFLAG tag, whereas the second was located downstream of a GFP sequence. The FLP recombinase was introduced into these lines through genetic crosses, enabling excision of the intervening endogenous genomic sequence. Five successive crosses were performed to establish the desired genotypes, as outlined in **Supplementary Figure 20**. Following recombination, expression of the FLAG-tagged CTCF orthologs was driven by the endogenous *CTCF* promoter, with the transgene-associated transcript retaining the endogenous 3′ untranslated region (3′ UTR).

Balancer chromosomes were used throughout the crossing schemes to maintain the desired genotypes and facilitate the identification of progeny. These chromosomes suppress recombination with their homologues and carry dominant visible markers that allow carriers to be distinguished at different developmental stages. The balancers employed included Bar, which produces reduced eye size; *ebony*, associated with dark body pigmentation; TM6 (Third multiple 6), used as a balancer chromosome; and *Ultrabithorax* (*Ubx*), which produces additional bristles on the halteres. The fly stocks used in this study, together with their corresponding descriptions, are listed in **Supplementary Table 2**.

### Protein Extraction and Western Blot

For protein extraction from mESCs, cell pellets were resuspended in 100 μL of urea buffer containing 8 M urea and 0.1 M Tris-HCl (pH 8.0), followed by vortexing. Samples were subsequently sonicated using a Bioruptor sonication system (Diagenode) at 4°C, with four cycles at high intensity, each consisting of 15 s of sonication followed by 45 s without sonication.

For *Drosophila* larval protein extraction, ten larvae were collected per sample and homogenized on ice in 100 μL of ice-cold cell lysis buffer comprising 5 mM HEPES (pH 8.0), 85 mM KCl, 0.5% NP-40, and protease inhibitors at a 1:25 dilution. The homogenates were centrifuged at 13,000 rpm for 5 min at 4°C to pellet the nuclei. The resulting pellets were resuspended in 30–50 μL of chilled nuclear lysis buffer containing 50 mM HEPES (pH 8.0), 10 mM EDTA, 0.5% NP-40, and protease inhibitors at a 1:25 dilution. The suspensions were then incubated at room temperature for 20 min. Protein concentrations were measured using either the Qubit Protein Assay or the Qubit Protein Broad Range Assay (Thermo Fisher Scientific), as appropriate for the sample concentration.

For mESC samples, 5 μg of nuclear protein extract was combined with 5× SDS loading buffer and heated at 95°C for 5 min. For *Drosophila* larval samples, 15 μg of nuclear extract was prepared in the same manner. The 5× SDS loading buffer was prepared in a final volume of 15 mL and contained 3 mL of 20% SDS, 3.75 mL of 1 M Tris-HCl (pH 6.8), 9 mg bromophenol blue, 1.16 mg DTT, and 4.5 mL glycerol, with double-distilled water added to the final volume. Protein samples were resolved by electrophoresis on NuPAGE 4–12% Bis-Tris gels (Invitrogen) at 120 V for 60–90 min. Proteins were subsequently transferred onto PVDF membranes using the Trans-Blot Turbo Transfer system at 25 V for 30 min. Membranes were blocked for 1 h at room temperature in 2.5% milk prepared in TBST (1× Tris-buffered saline containing 0.1% Triton X-100). Primary antibody incubation was carried out overnight at 4°C. The following day, membranes were washed three times for 5 min each with 0.1% TBST at room temperature and incubated with HRP-conjugated secondary antibodies diluted in 2.5% blocking milk for 1 h at room temperature. After three additional washes, chemiluminescent signals were detected using the Amersham Imager system. Antibodies used in this study are listed in **Supplementary Table 3**.

The Western blot image from Figure 3B has been cropped to eliminate an unrelated lane between the fly CTCF and mouse CTCF. The original image can be found in **Supplementary Figure 21**.

### Hi-C library preparation

For each replicate, 1 × 10⁶ mESCs were fixed in 2% formaldehyde prepared in 10% FCS/1× PBS for 10 min under gentle rotation. Fixation was stopped by adding 1.425 M glycine. The resulting cell pellet was resuspended in freshly prepared, ice-cold lysis buffer containing 50 mM Tris-HCl (pH 7.5), 150 mM NaCl, 5 mM EDTA, 0.5% IGEPAL CA-630, 1.15% Triton X-100, and 1× protease inhibitors. Samples were incubated on ice for 30 min and subsequently washed with 1× PBS. The resulting pellets were either snap-frozen for storage or processed immediately for downstream applications.

For fly material replicate, CNS were dissected from third-instar *Drosophila* larvae and collected in ice-cold 1× PBS. The collected material was centrifuged at 2,000 rpm for 5 min at 4°C, and the resulting pellet was resuspended in 500 μL of 10% FCS in PBS. The tissue was homogenized using a micropestle, followed by the addition of 250 μL of 4% formaldehyde prepared in 10% FCS/PBS. Samples were fixed for 10 min at room temperature under gentle rotation. Fixation was terminated by adding 100 μL of 1.425 M glycine, followed by gentle mixing through inversion on ice. The samples were then centrifuged at 4,000 × g for 8 min at 4°C, and the supernatant was discarded. The fixed pellet was resuspended in 200 μL of ice-cold cell lysis buffer containing 5 mM HEPES (pH 8.0), 85 mM KCl, 0.5% NP-40, and protease inhibitors at a 1:25 dilution. The material was gently homogenized on ice using a micropestle until the suspension became clear. The sample was subsequently centrifuged at 10,000 × g for 5 min at 4°C, and the supernatant was discarded. The pellet was resuspended in 500 μL of ice-cold cell lysis buffer and centrifuged again at 10,000 × g for 2 min at 4°C. After removal of the supernatant, the final pellet was snap-frozen until further processing.

Low-input Hi-C protocol was performed from fixed, lysed and snapped frozen cells as previously described^77^, with some minor modifications. Library amplification was carried out in four separate reactions of 12.5 μL each to optimize the number of PCR cycles and minimize over-amplification. Following amplification, a double-sided size selection was performed using AMPure XP beads. Library concentration was determined using the Qubit dsDNA High Sensitivity (HS) Assay Kit, while library fragment size distribution and overall quality were evaluated using a TapeStation system

### Hi-C data processing

Raw data was processed and filtered with Juicer using default parameters. The reference assemblies used were mm10 (mouse), dm6 (fruit fly), kPetMar1 (lamprey) and HT 2019 (Ciona robusta). For downstream analyses, Knight-Ruiz (KR) normalized Hi-C matrices in hic format at different resolutions were loaded into FAN-C. For the visualization of Hi-C heatmaps, hic files were loaded into Coolbox^78^.

### AB Compartment analysis

AB compartment analysis was performed using FAN-C in each replicate and chromosome individually in KR normalized matrices. After a high Pearson correlation between replicates was confirmed, the matrices of both replicates were merged. The first eigenvector was calculated again in the merged matrices, and the sign of the eigenvector was corrected if needed per individual chromosome depending on gene density (sea squirt, fly and lamprey) or H3K27ac ChIP-seq data when available^79^ (in mouse) signal in each chromosome independently. Chromosomes X and Y were excluded from this analysis in mouse.

### Insulation and chromatin domain analysis

Insulation scores (IS) were calculated from 10kb KR normalized matrices using FAN-C (window sizes: mouse = 250 kb, fly = 50 kb, lamprey = 50 kb to 1 Mb, Ciona robusta = 50 kb to 1 Mb). In mouse and fly replacement experiments, IS values were calculated in individual replicates to perform Pearson correlations. After confirming high Pearson correlations between replicates, IS values were recalculated from 10kb KR matrices with the replicates merged. Genome wide visualizations of IS values at genomic features of interest were obtained using the EnrichedHeatmap R package^80^. Boundary locations with their associated boundary scores (BS) were then calculated from IS values also with FAN-C. In the case of Ciona and lamprey, only boundaries that appeared recurrently using at least two different window sizes were kept for downstream analyses. For instance, chromatin domains in Ciona and lamprey were calculated as the intervening loci between recurrent boundaries. In lampreys, the presence and orientation of individual instances of CTCF motifs around TAD boundaries was predicted and quantified using FIMO^81^.

### Chromatin loops

Chromatin loops were calculated in mouse replacement experiments and in lamprey using Mustache with default parameters from 10kb KR normalized Hi-C matrices. Then, aggregate loop analyses across collections of relevant pairs of loci (i.e. loop anchors or CTCF binding site locations) were obtained using the R package mariner^82^. The orientations of the best CTCF motif instances obtained with FIMO for each loop anchor were kept to test the convergence rule in the lamprey Hi-C data.

### ChIP-seq

For mESC, 250000 cells per replicate were fixed with 1% formaldehyde in 10% FCS/1X PBS for 10 minutes rolling, then quenched by adding 1.425 M glycine. After a wash with 1X PBS, the pellet was either snap-frozen for storage or directly used for the downstream experiments. For fly embryonic material, brain tissue was processed as described for Hi-C experiments.

ChIP-seq experiments were performed using the Diagenode True MicroChIP kit with slight modifications. When using fly material, it was mechanically homogenised in 700 μL of RIPA buffer (140 mM NaCl, 8 mM NaHPO4, 2 mM NaH2PO4, 1% NP-40, 0.5% sodium deoxycholate, 0.05% SDS, 0.2% PMSF) supplemented with protease inhibitors, then centrifuged at 16000 g for 5 minutes at 4°C. 250000 cells per replicate were used for subsequent lysis and sonication using the Diagenode Bioruptor Sonication device (HIGH: 30sec ON, 30sec OFF for 15 cycles at 4°C). mESC lysates were sonicated with the same settings for 20 cycles. For spike-in experiments, 5 ng of spike-in chromatin (Active Motif) and 1 μg of spike-in antibody (Active Motif) were added before the overnight incubations with the target-specific antibody. For the 3XFLAG pull-down, 2 μg of Sigma-Aldrich anti-FLAG M2 antibody was used. For cohesin pull-down, 2 μg of Abcam anti-Rad21 antibody was used. ChIP-seq analysis

ChIP-seq was performed using the Diagenode True MicroChIP kit, with minor modifications. For *Drosophila* material, the tissue was mechanically homogenized in 700 μL of RIPA buffer containing 140 mM NaCl, 8 mM NaHPO₄, 2 mM NaH₂PO₄, 1% NP-40, 0.5% sodium deoxycholate, 0.05% SDS, and 0.2% PMSF, supplemented with protease inhibitors. The homogenate was centrifuged at 16,000 × g for 5 min at 4°C. Chromatin from 2.5 × 10⁵ cells per replicate was subsequently lysed and sonicated using a Bioruptor sonication system (Diagenode). Sonication was performed at high intensity at 4°C, with 30 s ON and 30 s OFF intervals. Fifteen cycles were applied to *Drosophila* samples, whereas mESC samples underwent 20 cycles under the same conditions.

For spike-in normalization experiments, 5 ng of spike-in chromatin and 1 μg of the corresponding spike-in antibody (Active Motif) were added before overnight incubation with the target-specific antibody. For immunoprecipitation of 3×FLAG-tagged proteins, 2 μg of anti-FLAG M2 antibody (Sigma-Aldrich) was used. For cohesin-associated chromatin enrichment, immunoprecipitation was performed using 2 μg of anti-Rad21 antibody (Abcam).

### ChIP-seq data analysis

CTCF-FLAG ChIP-seq reads were first trimmed with Trim Galore (https://github.com/felixkrueger/trimgalore). Trimmed reads were then aligned to either the mm10 (mouse replacements) or the dm6 (fly replacements) genome assemblies using bowtie2^83^ and sam alignments were converted to bam, sorted and deduplicated using sambamba^84^. Drosophila spike-in reads in mouse samples were detected and quantified by repeating the previous steps using the dm6 assembly of flies. Scaling factors were then calculated and used as an input for the calculation of bigwig files from deduplicated bam files using deeptools^85^ bamCoverage (parameters: --extendReads --centerReads –scaleFactor <SCALE factor>). In the case of fly experiments, RPGC normalization was used in deeptools bamCoverage (--normalizeUsing RPGC --effectiveGenomeSize <GENOME size>). ChIP-seq peaks were then calculated using macs2^86^ callpeak in the different replicates separately and also in the two replicates combined. These three sets of peaks per experiment were used as an input to IDR^87^ to obtain a conservative set of reproducible CTCF peaks following the ENCODE recommendations. Normalized ChIP-seq signals were then visualized over CTCF peaks using the EnrichedHeatmap R package and the normalized signal was used to cluster those CTCF sites according to their binding dynamics using k-means. De-novo motif prediction in the different clusters was performed with STREME^88^ using the R packages memes^89^ as an interface and for the visualization of logos. ChIPseeker was used to characterize the genomic location of ChIP-seq peaks (i.e. distal, intronic, promoter). Visualizations of ChIP-seq^90^ signal on individual loci combined with Hi-C were obtained with Coolbox^78^.

### Methylation data analyses

CpG methylation calls obtained from Whole Genome Bisulfite Sequencing (WGBS) data in mESCs were downloaded from GEO (GSE248933) in bedgraph format^91^. The bedgraph was split by strand and represented alongside ChIP-seq and conservation data using the EnrichedHeatmap R package. In the zoom out plot, methylation levels in non-CpG positions were imputed as the mean of the methylation of the two flanking CpGs.

### RNA-seq

Total RNA was extracted from 1 × 10⁶ mESCs per replicate. Cell pellets were resuspended in TRIzol reagent, and RNA purification was performed using the Direct-zol RNA MiniPrep kit (Zymo Research), following the manufacturer’s instructions. A total of 500 ng of purified RNA was used as input for mRNA enrichment with the NEBNext Poly(A) mRNA Magnetic Isolation kit (New England Biolabs).

RNA-seq libraries were subsequently prepared using the NEBNext Ultra II Directional RNA Library Prep Kit (New England Biolabs), according to the manufacturer’s protocol. Library quality and fragment size distribution were assessed using a TapeStation system.

### RNA-seq data analysis

Alignment and gene expression quantification were performed using the PiGx^92^ RNA-seq pipeline with default parameters using the Mus_musculus.GRCm39 reference genome and annotation provided by Ensembl. Briefly, reads were trimmed with Trim Galore (https://github.com/FelixKrueger/TrimGalore) and aligned with STAR^93^. Per-gene quantification was performed using the GenomicAlignments::summarizeOverlaps R function^94^. Differential analysis was then performed using DESeq2^95^. Specifically, the FDR-corrected P values of all pairwise comparisons between genotypes were computed using DESeq2::contrast function (using Wald tests).

### 3’RACE

The 3′ end of the transcript was characterized using the 3′ RACE System for Rapid Amplification of cDNA Ends kit (Invitrogen), following the manufacturer’s instructions. Amplification was performed using the gene-specific primer 5′-ATCAGCAAACTACGCCGACATT-3′. The resulting PCR product was separated by gel electrophoresis, and the fragment of interest was excised and purified using the GeneJET Gel Extraction Kit (Thermo Fisher Scientific). The purified product was subsequently submitted to LGC Genomics for Sanger sequencing using the same gene-specific primer, according to the provider’s guidelines.

### Viability Assay

The fitness of homozygous and heterozygous CTCF ortholog-replacement mutants was evaluated by monitoring developmental progression and adult survival. For each genotype, three independent groups of 30–40 third-instar larvae were transferred to separate vials. The numbers of pupae and successfully eclosed adults were recorded over the 14-day period following larval hatching. The mean survival percentages at each developmental stage and the corresponding standard deviations were subsequently calculated. Adult survival was assessed using Kaplan–Meier survival analysis, which was used to estimate survival probabilities over time.

### Locomotion Assay

Locomotor behaviour in newly eclosed adult flies was assessed by recording their movement in lidded 6-well and 12-well plates. Videos lasting 10–30 s were acquired using a Leica MC120 microscope camera and Leica Application Suite software. Video recordings were subsequently analyzed in ImageJ to manually measure the total distance travelled by each individual. Locomotor speed was calculated by dividing the distance travelled (in millimetres) by the recording duration (in seconds), yielding a movement rate expressed in mm/s.

### Protein alignments

Sequences were aligned using MAFFT v7.490^96^ in its accurate mode (linsi) and alignments were visualized and exported using Jalview^97^.

### Phylogenetic analyses

Five sequences from reference species: *S. purpuratus* [SPU.012546.3a or XP_011671461.2B); *D.melanogaster* (Q9VS55); *C. robusta* (ENSCING00000006765); *D. rerio* ENSDARG00000056621, and *M. musculus* ENSMUSG00000005698 were aligned using mafft^98^ with its accurate mode, L-INS-i mode.

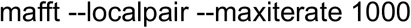

The region corresponding to the Zn-finger central regions was extracted, removing upstream and downstream regions. This alignment was used to generate a seed HMM profile using hmmer^99^ with hmmbuild. The profile was used to search sequences from a metazoan collection, MAteDB2^100^ that contains ∼1000 metazoan proteomes. hmmsearch was used with the following parameters:

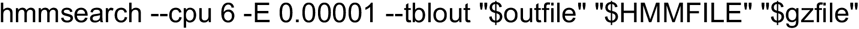

Accepted hits were combined with known/reference CTCF-like sequences into a representative alignment containing 83 sequences. A maximum-likelihood phylogeny of the trimmed central CTCF zinc-finger region was subsequently generated, based on a pruned 120-sequence alignment. Since this is a domain-region tree, not a full-length protein tree, it reflects phylogenetic signal in the zinc-finger region specifically. The tree runs were performed with IQ-TREE 1.6.12 (https://iqtree.github.io/) using ModelFinder, ancestral state reconstruction, ultrafast bootstrap, and -bnni. A total of 30,000 ultrafast bootstrap replicates were performed, with automatic model selection. Trees were visualized with iTOL^101^ and exported to pdf.

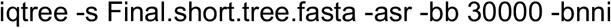

## Data availability

Datasets have been deposited in GEO repository (GSEXXXXXX).

## Code availability

The custom R and python code to reproduce the analyses and the figures is available in GitLab (https://gitlab.com/rdacemel/replace-ctcf).

