## Supplementary Figures for "Evolution of CTCF underlies the establishment of loop extrusion as a predominant chromatin folding mechanism in vertebrates"

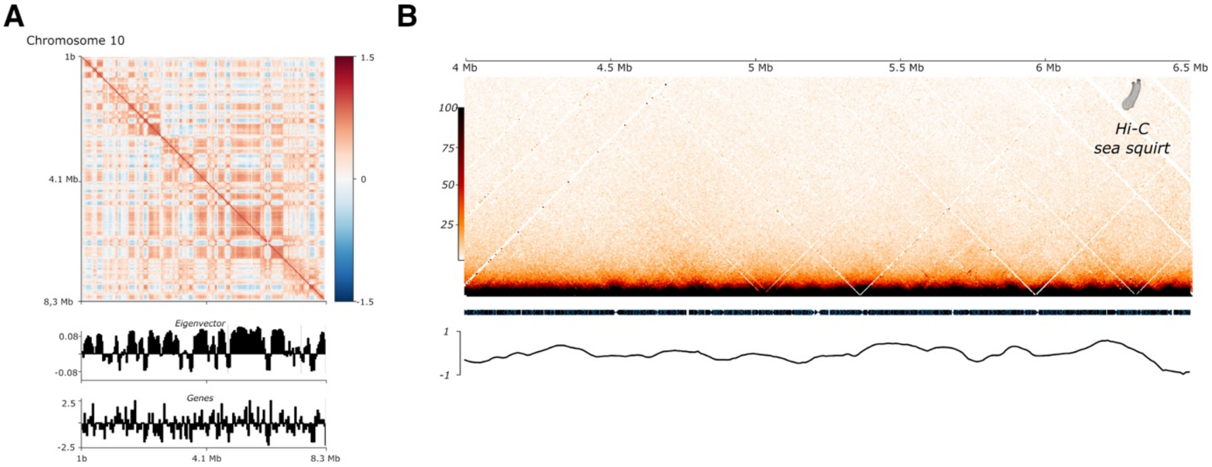


***Supplementary Figure 1. 3D chromatin organization in tunicates. A.*** *A/B compartmentalization in tailbud embryos from Ciona robusta. Representative example from chromosome 10 is shown (resolution = 40 kb). Similar results were obtained in other chromosomes. Top, Pearson matrices showing A/B compartmentalization. Bottom, eigenvector and gene density plots. Note the enrichment in gene density for A compartment (positive eigenvector signal). B. Hi-C maps from tailbud embryos of Ciona robusta. A representative locus is shown. Similar results were obtained in other genomic regions or when profiling heart or mesenchymal cells. Lower panel, Insulation scores for the same genomic region. Note the overall absence of TAD signal.*

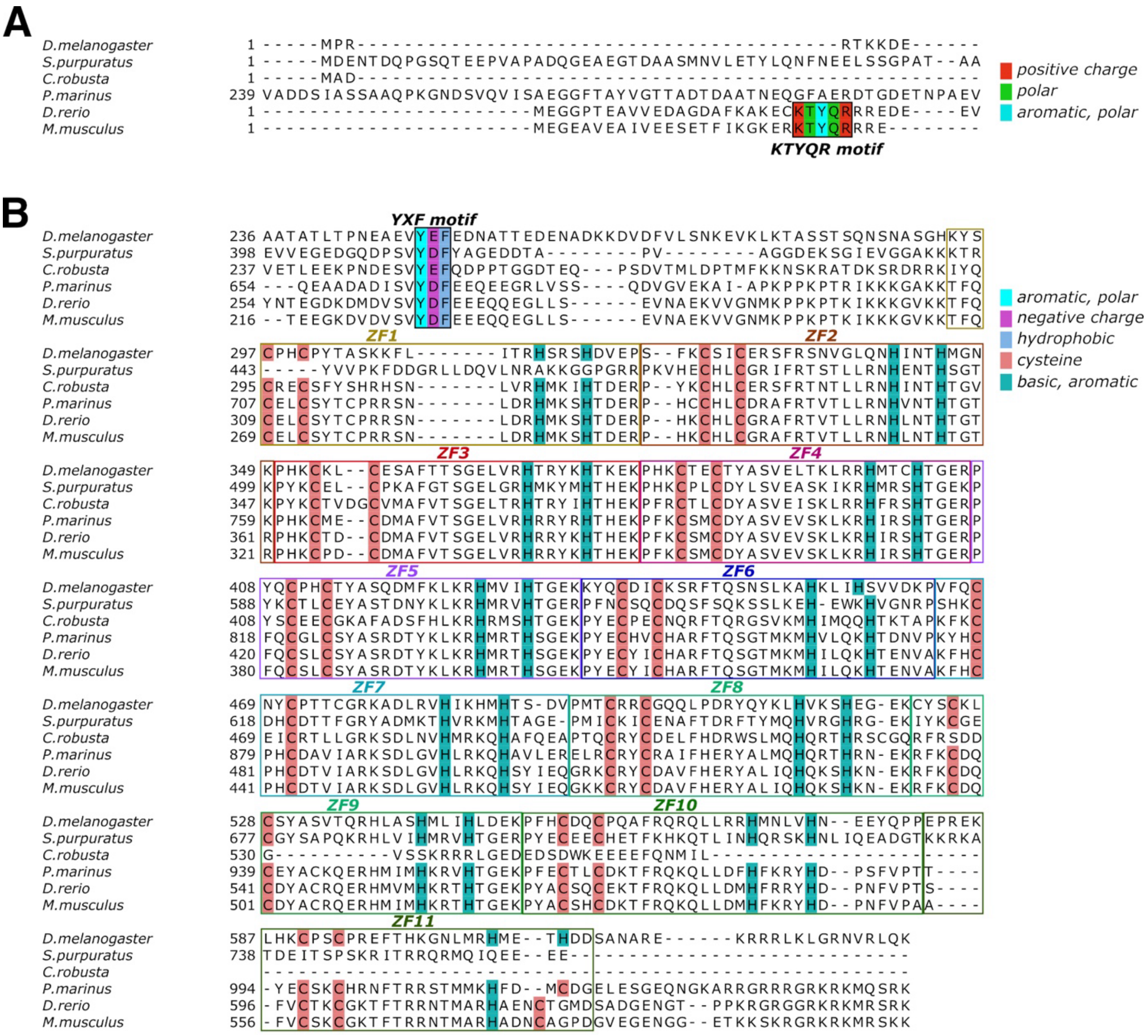


***Supplementary Figure 2. Evolution of the CTCF protein across bilateria. A**. ClustalW alignment of the N-terminal region containing the KTYQR motif. This motif is conserved only in gnathostomes (jawed vertebrates). **B.** ClustalW alignment of the N-terminal region containing the YXF motif, together with the zinc finger (ZF) domain. The YXF motif is conserved across Bilateria. Despite variation in the total number of ZFs, all species present ZF3–ZF7, which are critical for DNA binding. Cysteine (C) and histidine (H) residues, which coordinate zinc ions, are marked.*

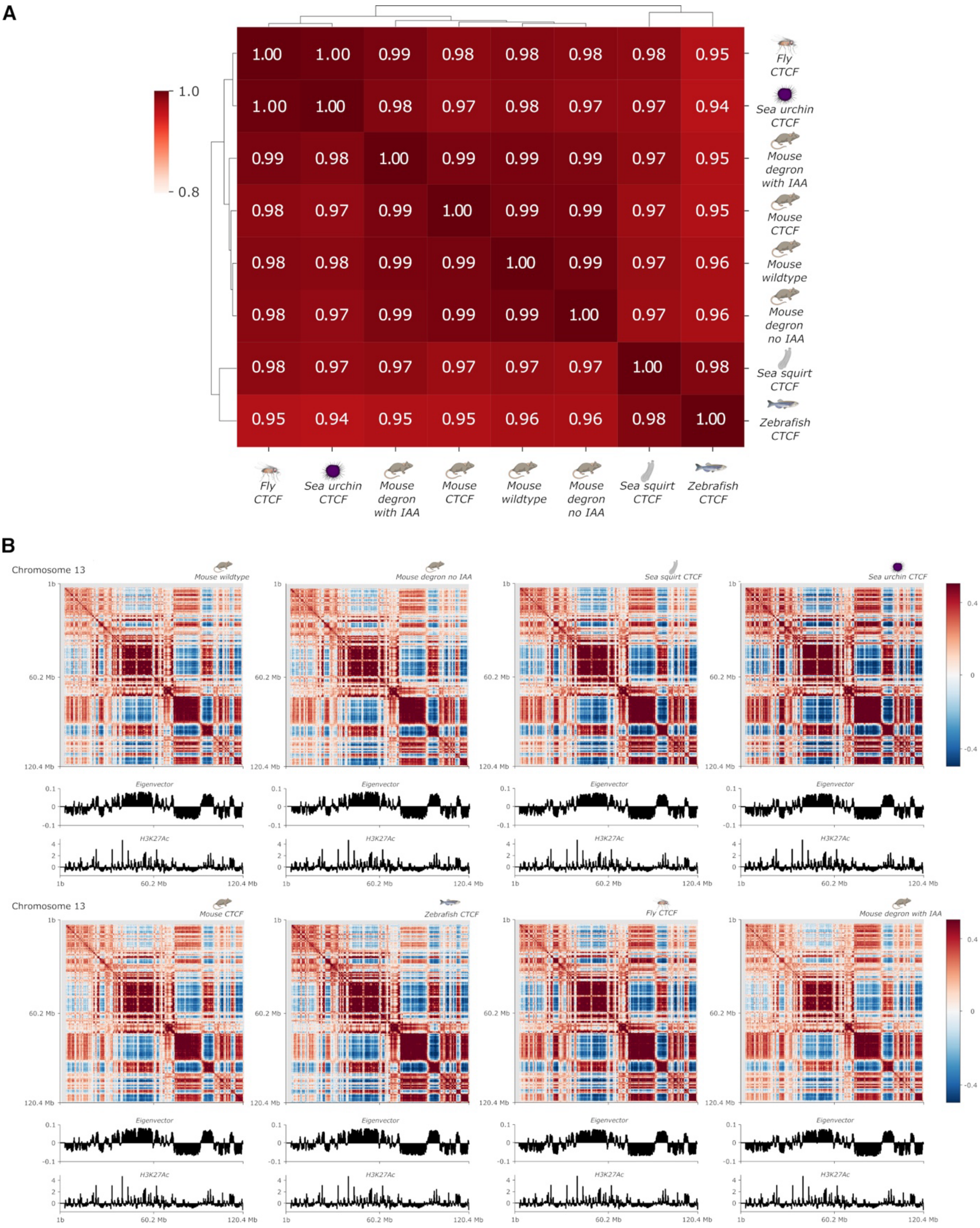


***Supplementary Figure 3. A/B compartmentalization in CTCF orthologous replacement mESC lines. A.*** *Correlation of eigenvector signal from Hi-C experiments. Note that all samples display a high degree of correlation.* ***B.*** *Top, Pearson matrices showing A/B compartmentalization of chromosomes (resolution = 250 kb). Bottom, eigenvector and H3K27Ac ChIP-seq plots. Note the enrichment in H3K27Ac signal for A compartment (positive eigenvector signal). Representative examples from chromosome 13. Similar results are obtained in other chromosomes*

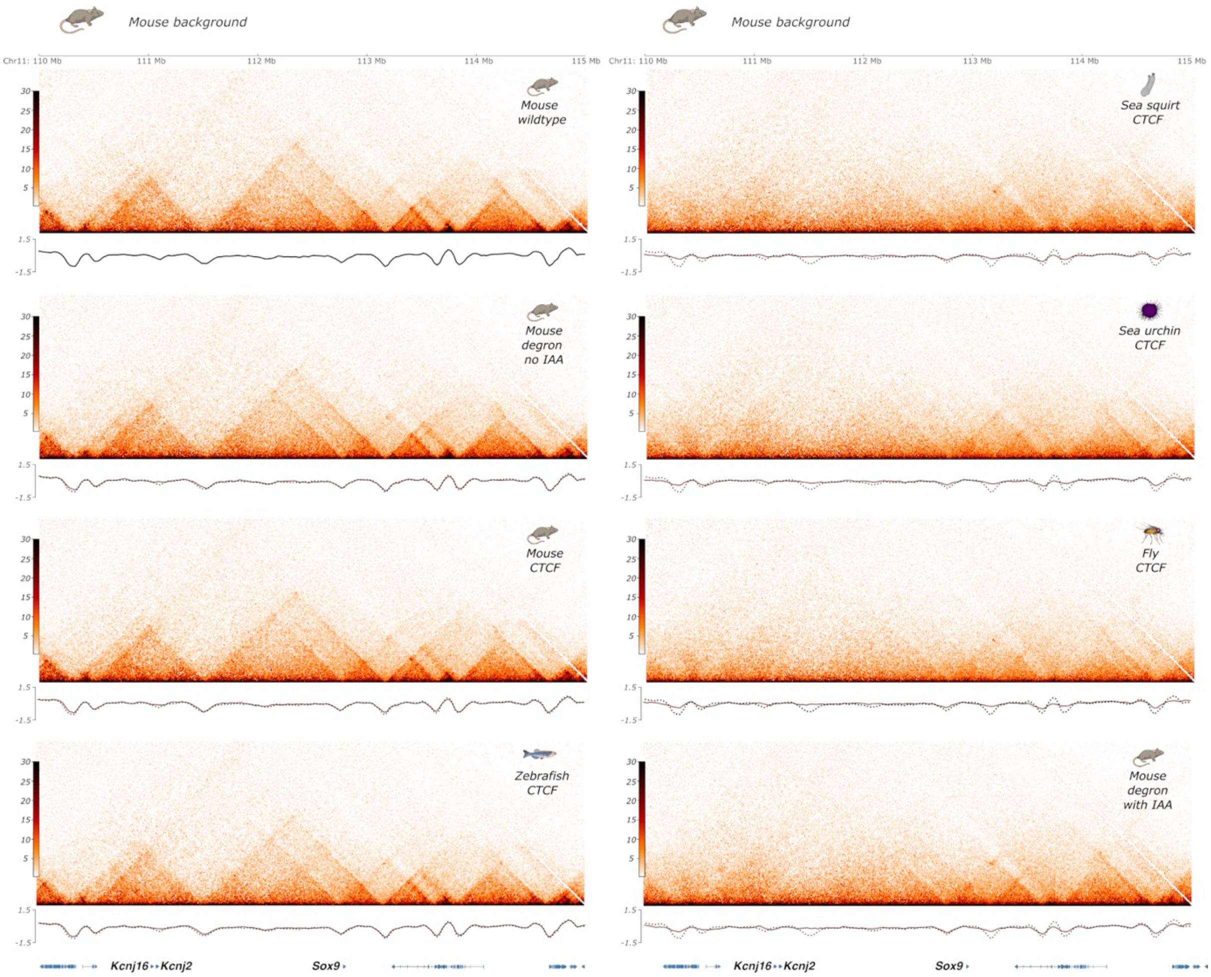


***Supplementary Figure 4. TAD organization in CTCF orthologous replacement mESC lines.*** *Hi-C maps of the Sox9 locus in different samples. Lower panel. Insulation scores for the same genomic region. Note the existence of two distinct groups of interaction profiles, containing vertebrate (left) and invertebrate (right) samples.*

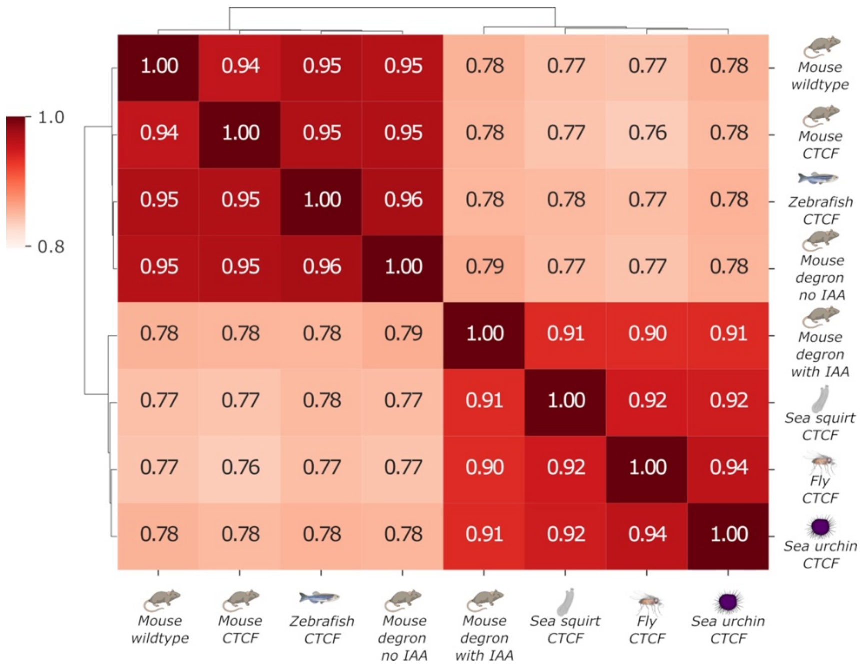


***Supplementary Figure 5. Correlation of insulation score signal from Hi-C experiments in mESC orthologous replacement lines.*** *Note the clustering of two large sample groups, containing invertebrate and vertebrate samples, respectively.*

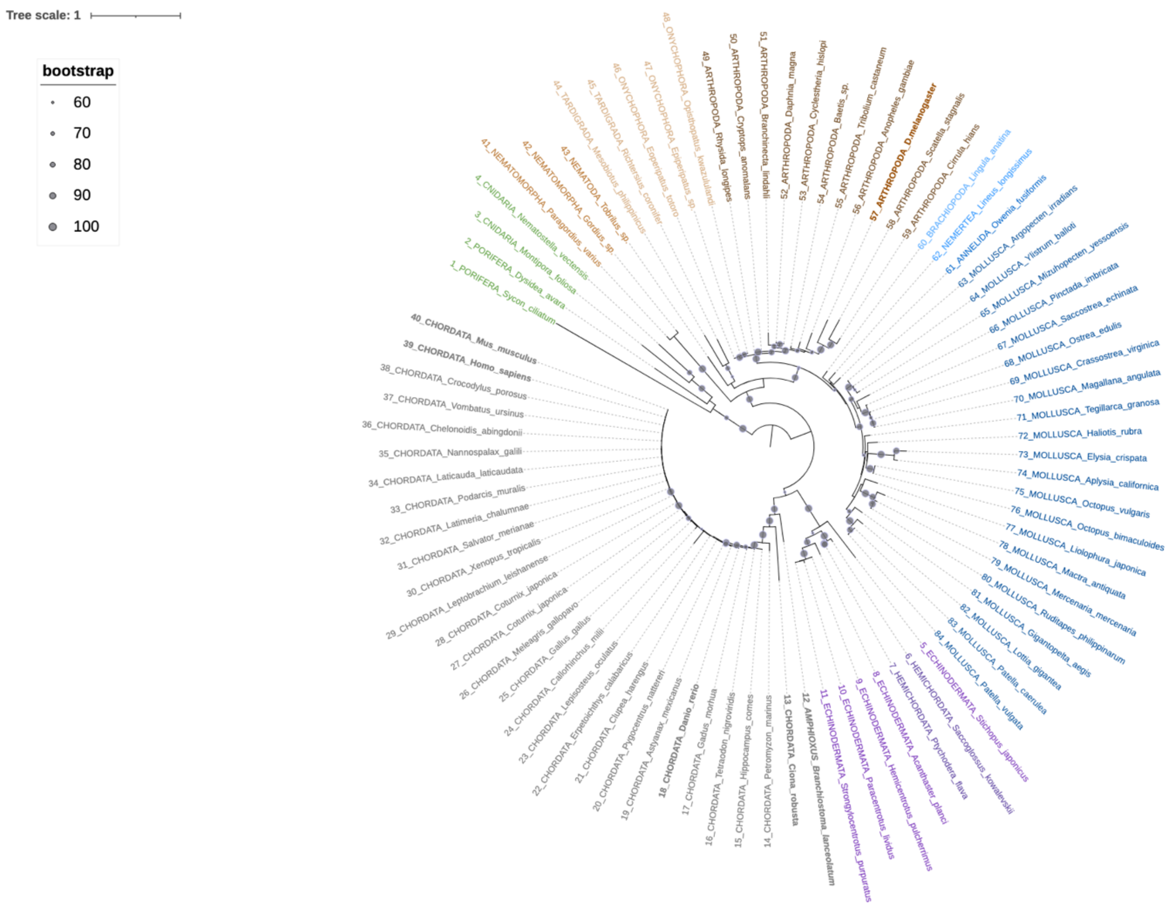


***Supplementary Figure 6. Maximum-likelihood phylogeny of the CTCF zinc-finger region (2-10) across bilateria.*** *Unrooted maximum-likelihood phylogeny of the central CTCF zinc-finger region (2-10 ZFs) for 84 representative taxa spanning major bilaterian lineages, including Echinodermata, Hemichordata, Chordata, Nematomorpha, Nematoda, Tardigrada, Onychophora, Arthropoda, Brachiopoda, Nemertea, Annelida and Mollusca. Porifera and Cnidaria represent outgroups. Tip labels are colour-coded according to phylum for clarity. Bold tip labels indicate relevant species. Gray circles at internal nodes represent bootstrap support values obtained from 30,000 bootstrap replicates, with circle size proportional to the level of support. Only bootstrap values greater than 60% are displayed. Branch lengths are proportional to evolutionary distance, with the corresponding scale indicated in the upper-left corner.*

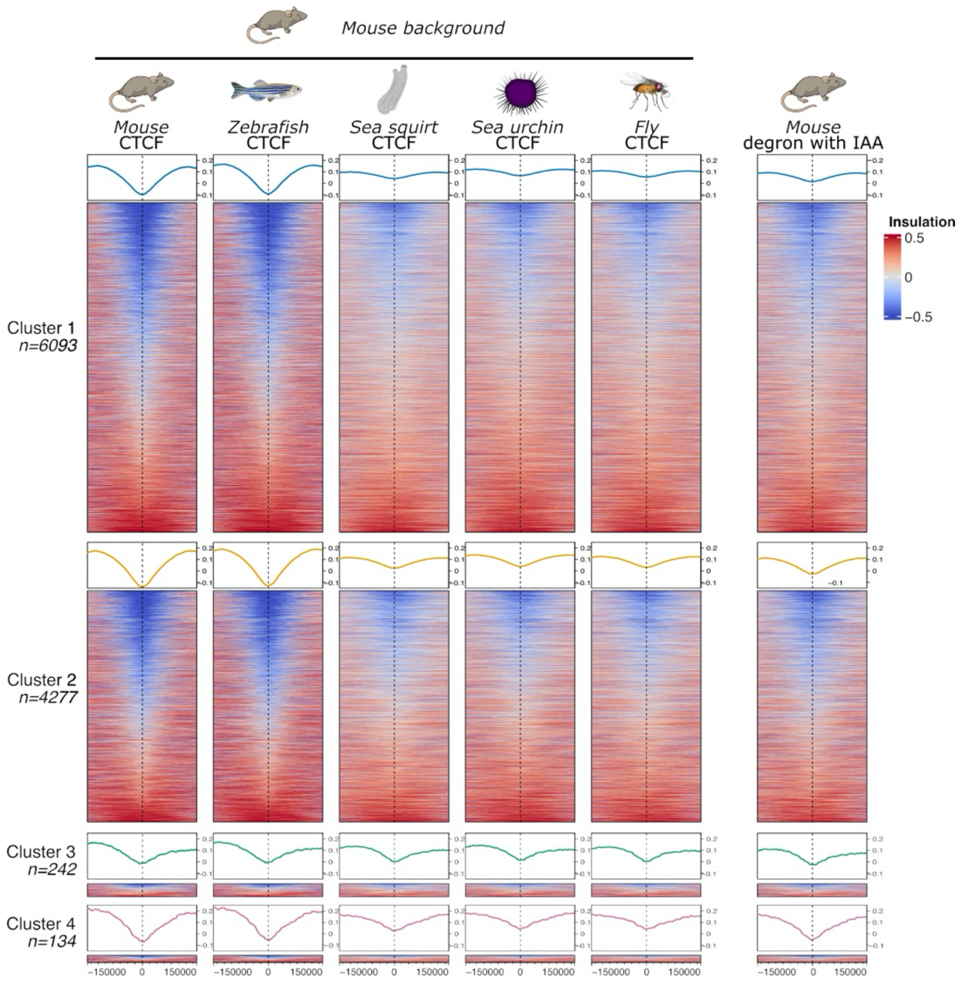


***Supplementary Figure 7. Boundary analyses in CTCF orthologous replacement mESC lines.*** *Insulation scores across genomic regions corresponding to the clusters of CTCF ChIP-seq signal shown in Figure 2C. Note that, despite chromatin binding, sea squirt CTCF exhibits similar insulation scores in clusters 2 and 3 to those of other invertebrate orthologs, which do not bind to these regions.*

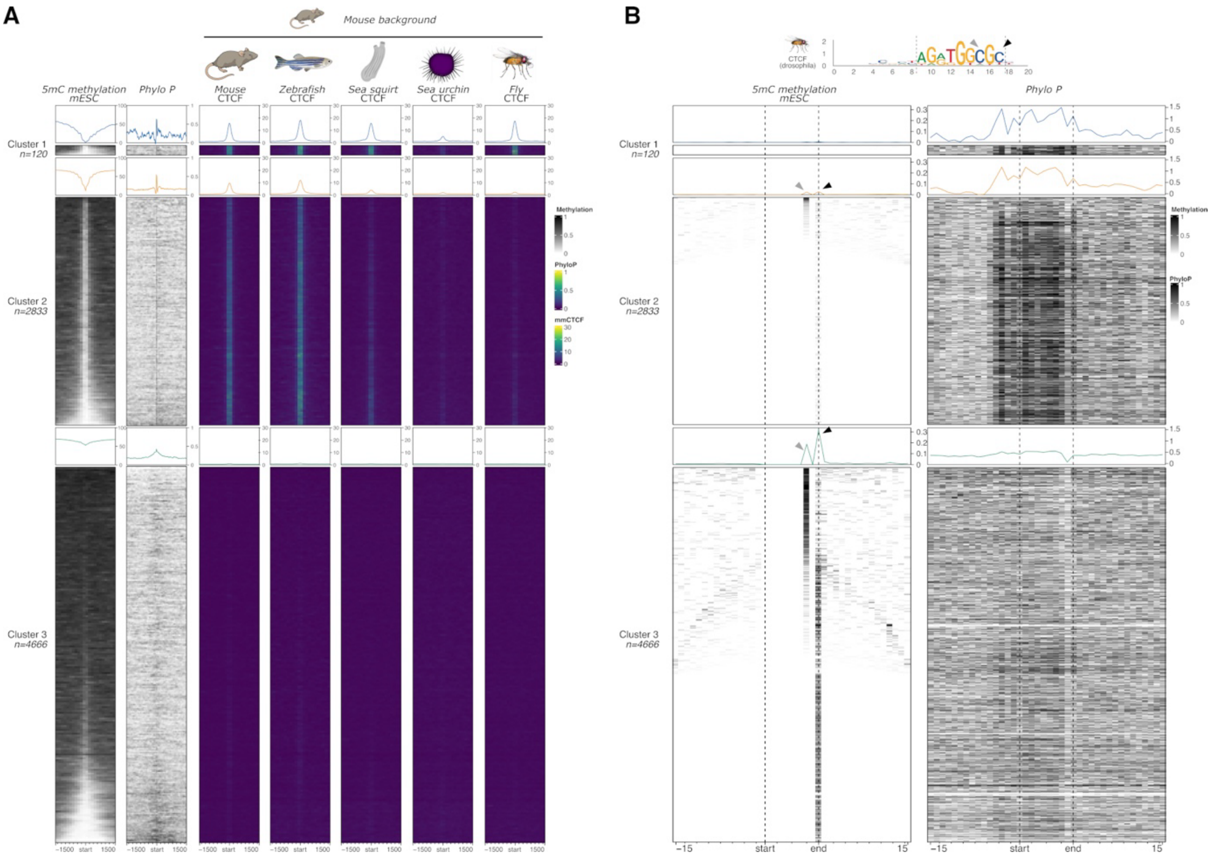


***Supplementary Figure 8. 5mC methylation analyses in mESC. A.*** *Clustering of CTCF ChIP-seq signal in regions containing the invertebrate (Drosophila) binding motif. Note that unbound regions (cluster 3) display a higher degree of 5mC methylation in mESC and tend to be less conserved.* ***B****. Zoomed region of the clustering shown in a. Dashed line represents the position of the invertebrate (Drosophila) CTCF binding motif. Arrows denote the cytosines that are susceptible to 5mC methylation. Note that the terminal cytosine (black arrow) shows higher levels of 5mC methylation, particularly in cluster 3.*

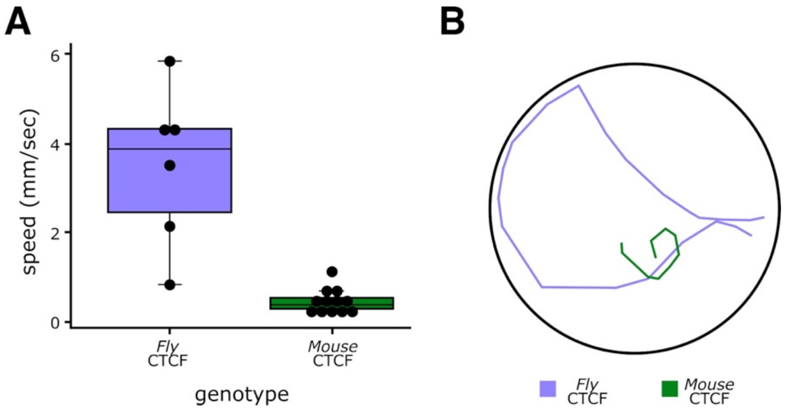


***Supplementary Figure 9: Locomotion assay of CTCF orthologous replacement fly lines. A.*** *Fly speed in mm/sec. One-way ANOVA with posthoc Tukey HSD. SD indicated by error bars, p-value < 0.0001.* ***B.*** *Representative paths followed by a single male animal of each replacement line*

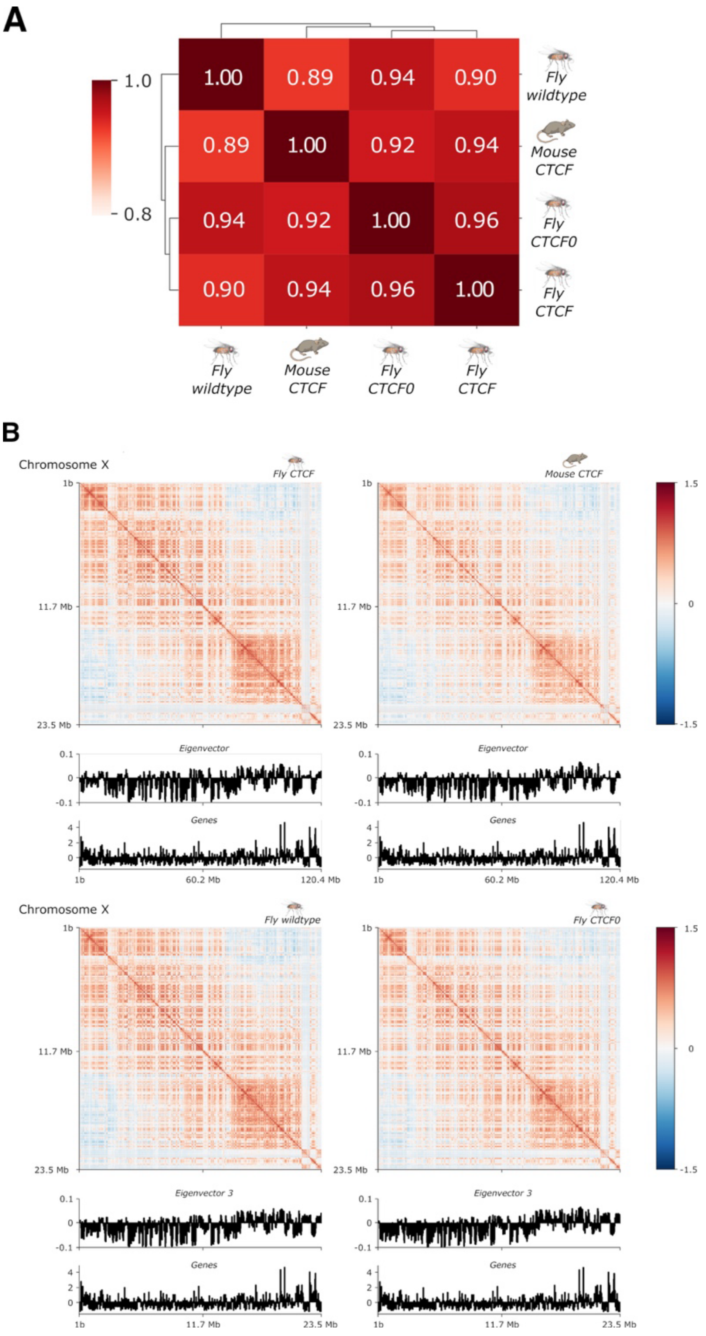


***Supplementary Figure 10. A/B compartmentalization in CTCF orthologous replacement in fly lines. A.*** *Correlation of eigenvector signal from Hi-C experiments. Note that all samples display a high degree of correlation.* ***B.*** *Representative examples from chromosome X. Similar results are obtained in other chromosomes. Top, Pearson matrices showing A/B compartmentalization of chromosomes (resolution = 40 kb). Bottom, eigenvector and gene density plots. Note the enrichment in gene density for A compartment (positive eigenvector signal).*

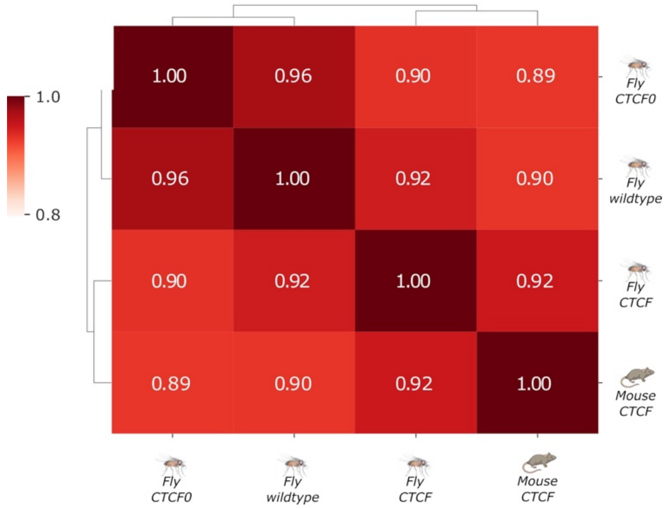


***Supplementary Figure 11. Correlation of insulation score signal from Hi-C experiments in fly orthologous replacement lines****. Note that all samples display a high degree of correlation.*

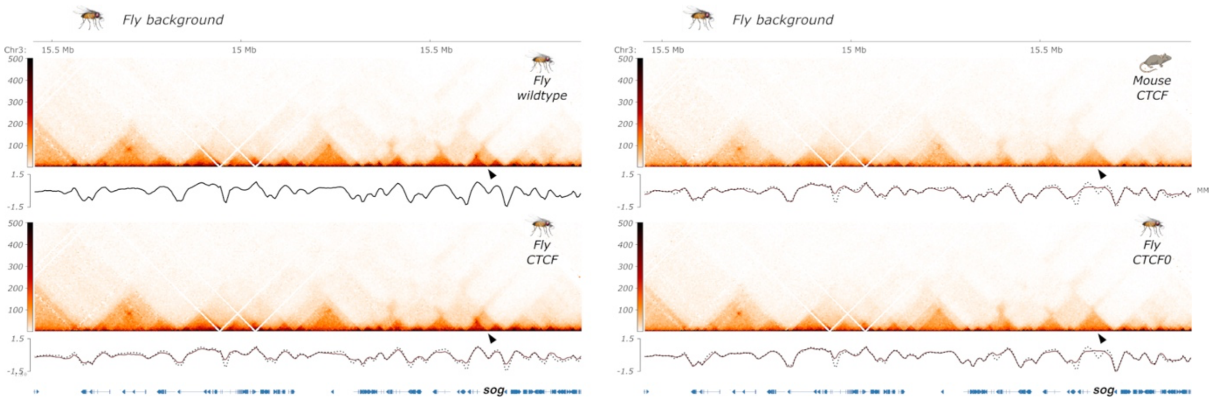


***Supplementary Figure 12. TAD organization in CTCF orthologous replacement in fly lines.*** *Hi-C maps from CNS of the sog locus in different samples. Lower panel. Insulation scores for the same genomic region. Note the presence of two distinct groups of interaction profiles, containing wildtype and fly CTCF replacements (left) and another containing mouse CTCF replacement and fly CTCF0 (right) samples. Black arrow denotes a differential TAD boundary between both groups.*

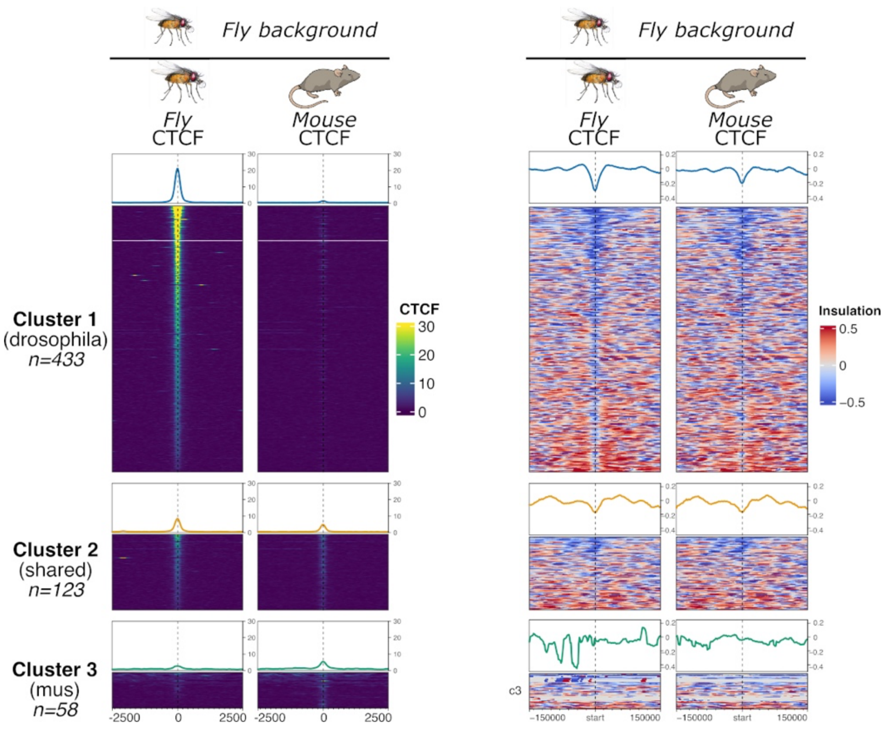


***Supplementary Figure 13. Insulation associated with CTCF orthologous binding in fly replacement lines.*** *Left, clustering of CTCF ChIP-seq signal at specific genomic regions. Right, insulation signal for the same clusters. Note the decrease in insulation in mouse CTCF samples for cluster 1.*

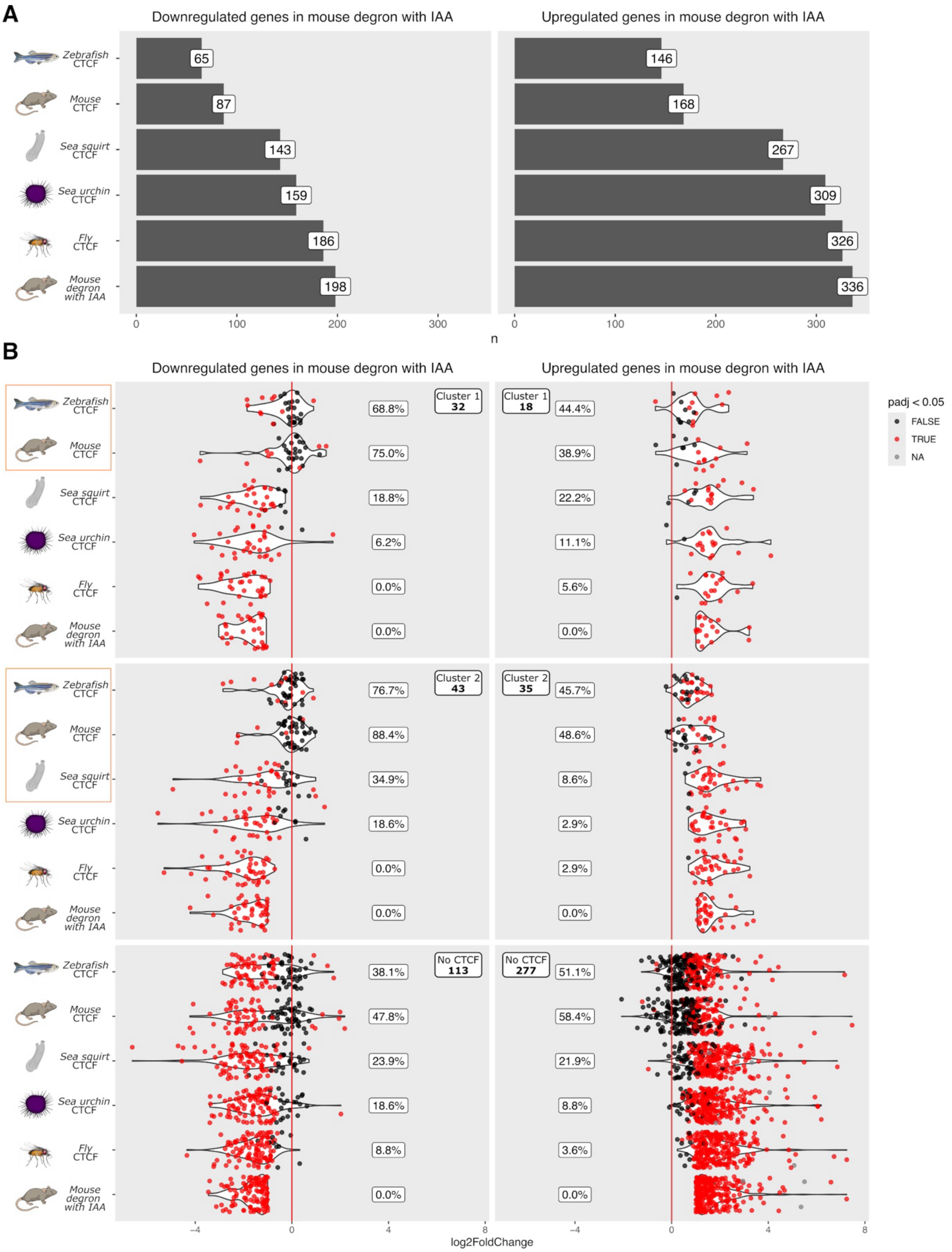


***Supplementary Figure 14. RNA-seq analyses in CTCF orthologous replacement in mESC lines. A.*** *Number of dysregulated genes in mESC lines.* ***B.*** *Percentage of rescued genes associated with promoter-proximal CTCF binding. Clusters relate to those shown for CTCF ChIP-seq binding analyses from Figure 2B (clusters 3 and 4 contain no CTCF-bound genes). The number of genes per category is indicated in the upper corner of each plot. Orange rectangles indicate those orthologous replacements where CTCF is bound at the gene promoters.*

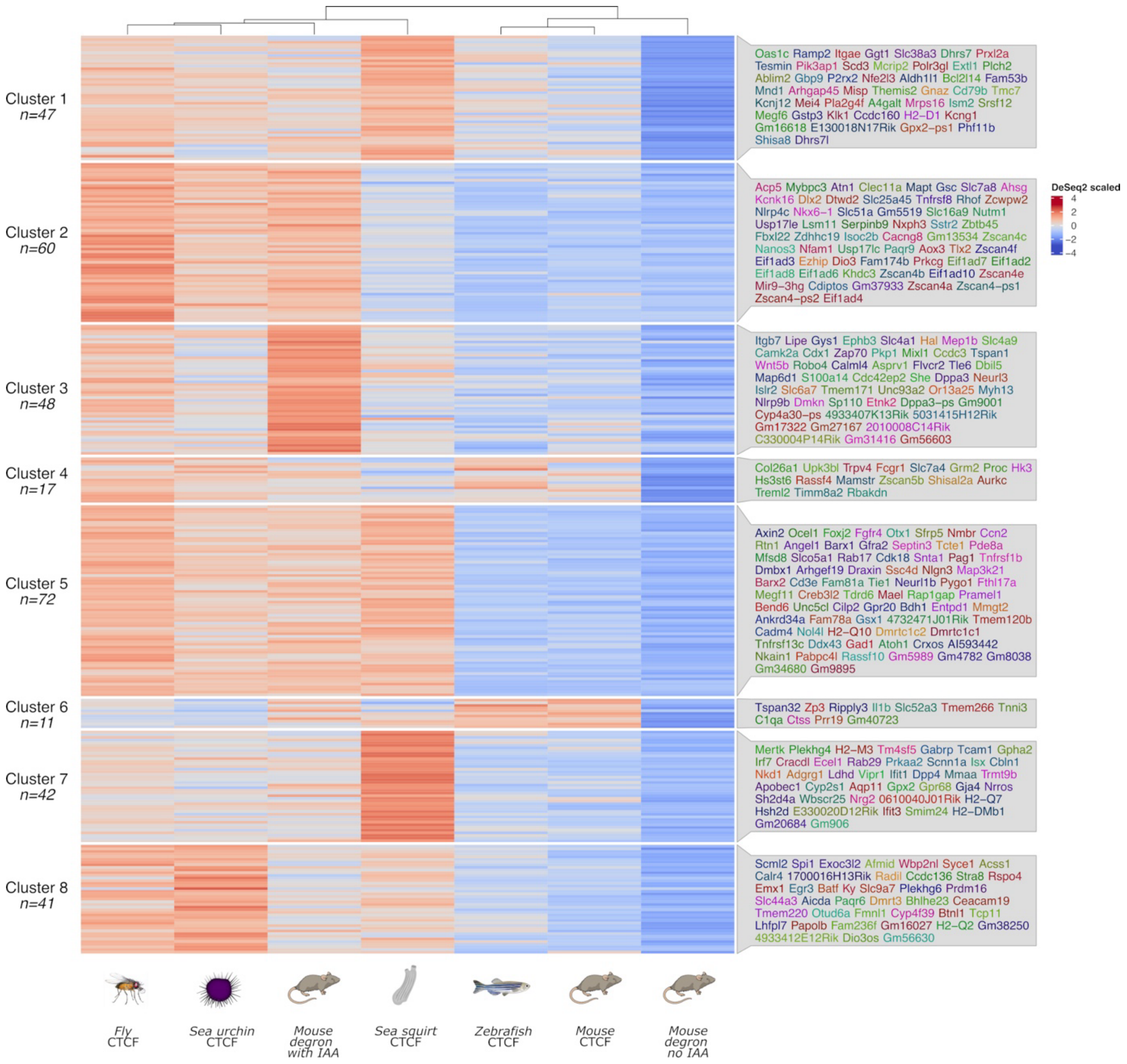


***Supplementary Figure 15. Upregulated genes in CTCF orthologous replacement mESC lines.*** *Cluster 2 represents genes that are rescued upon replacement with chordate CTCF orthologs (sea squirt, zebrafish or mouse).*

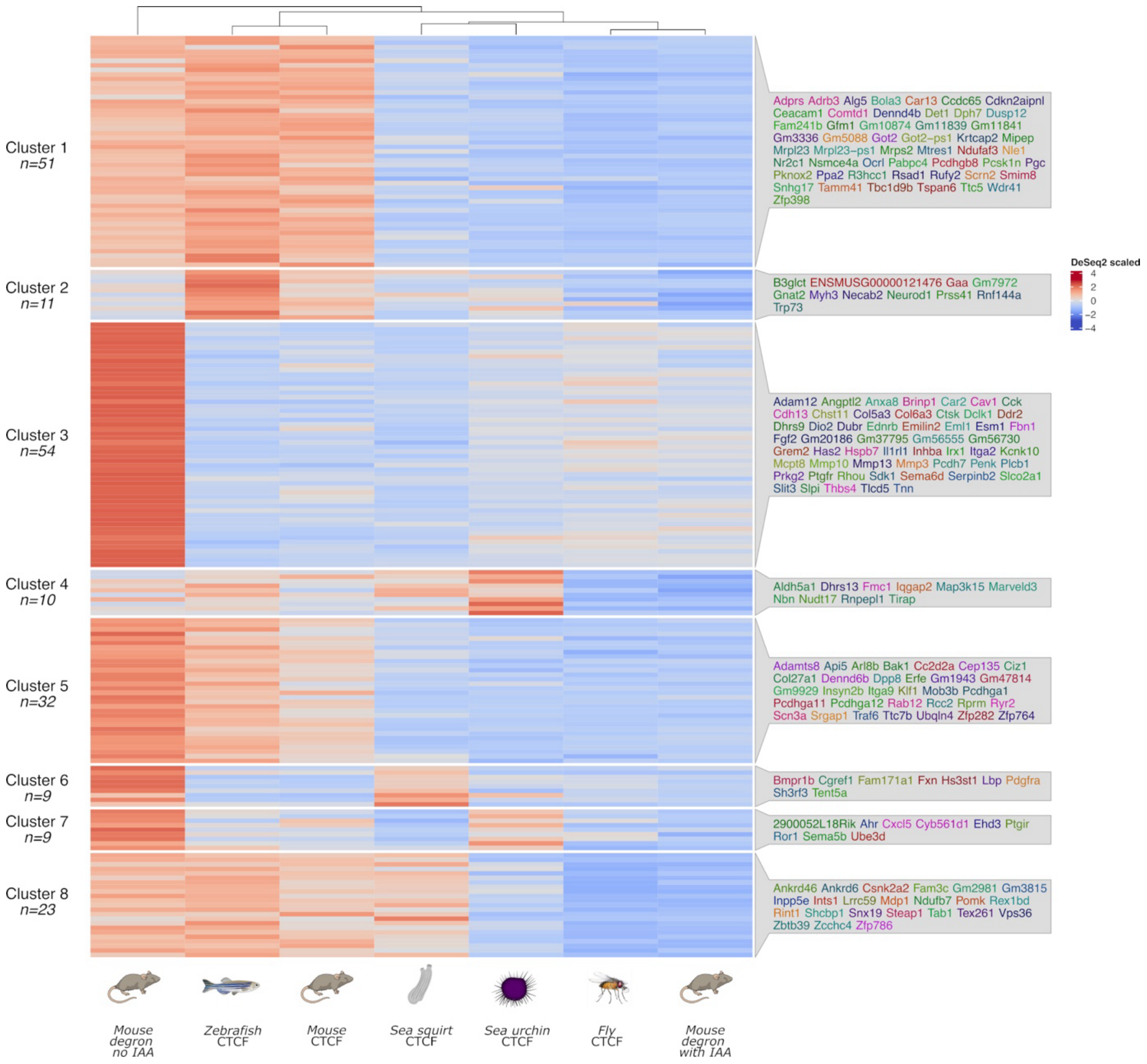


***Supplementary Figure 16. Downregulated genes in CTCF orthologous replacement mESC lines.*** *Cluster 8 represents genes that are rescued upon replacement with chordate CTCF orthologs (sea squirt, zebrafish or mouse).*

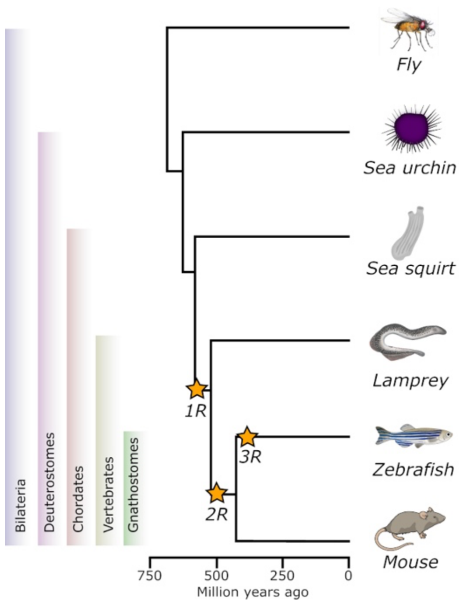


***Supplementary Figure 17. Evolutionary tree of representative bilaterian species.*** *Evolutionary tree showing divergence time between species used in this study. Orange stars represent each round of vertebrate-specific Whole Genome Duplications (1R, first round; 2R, second round).*

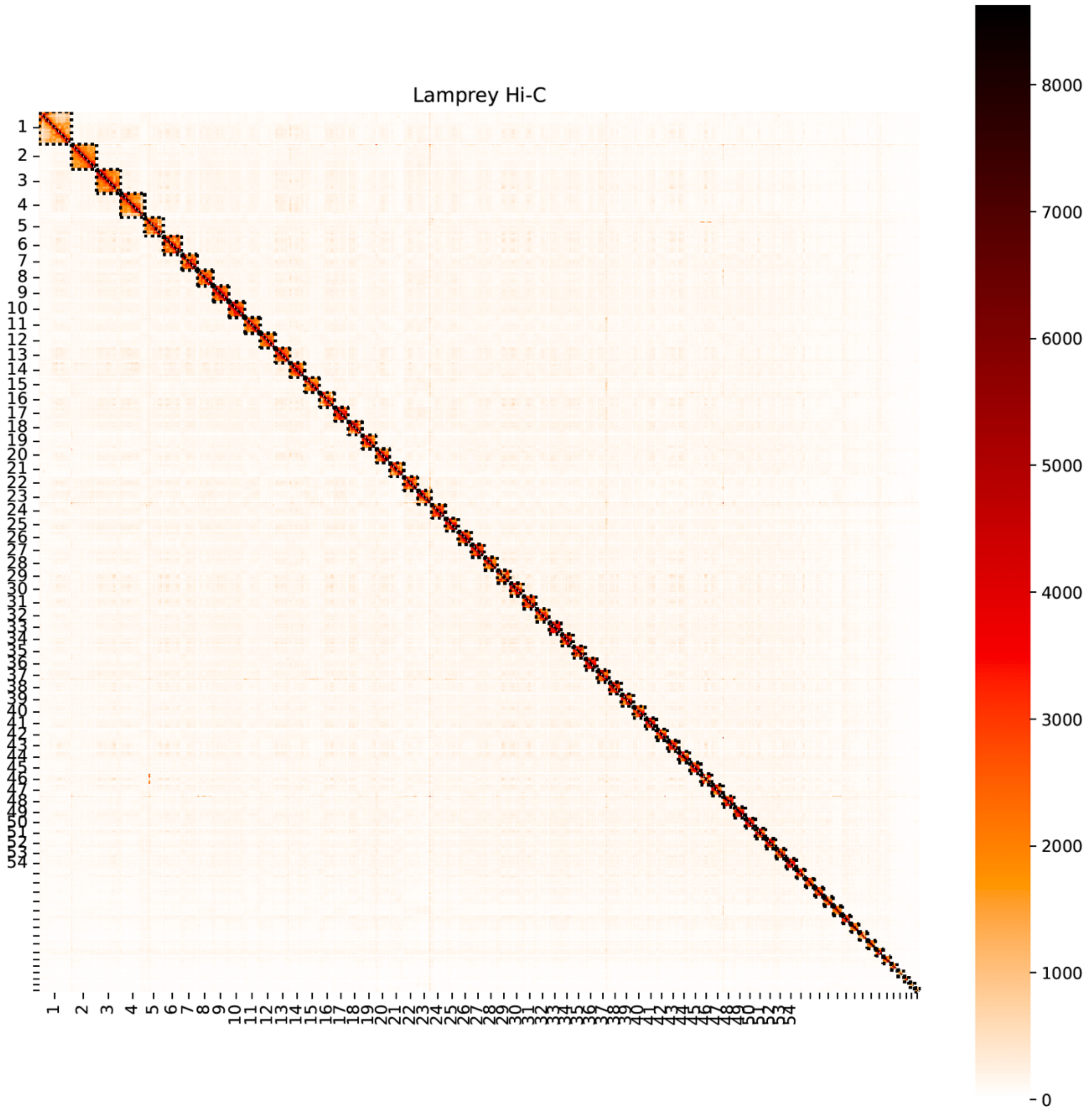


***Supplementary Figure 18. Chromosome territories in cyclostomes.*** *Hi-C map at 1Mb resolution of the entire sea lamprey (Petromyzum marinus) genome. Chromosome ends are delineated with dashed lines. Intrachromosomal contacts are enriched.*

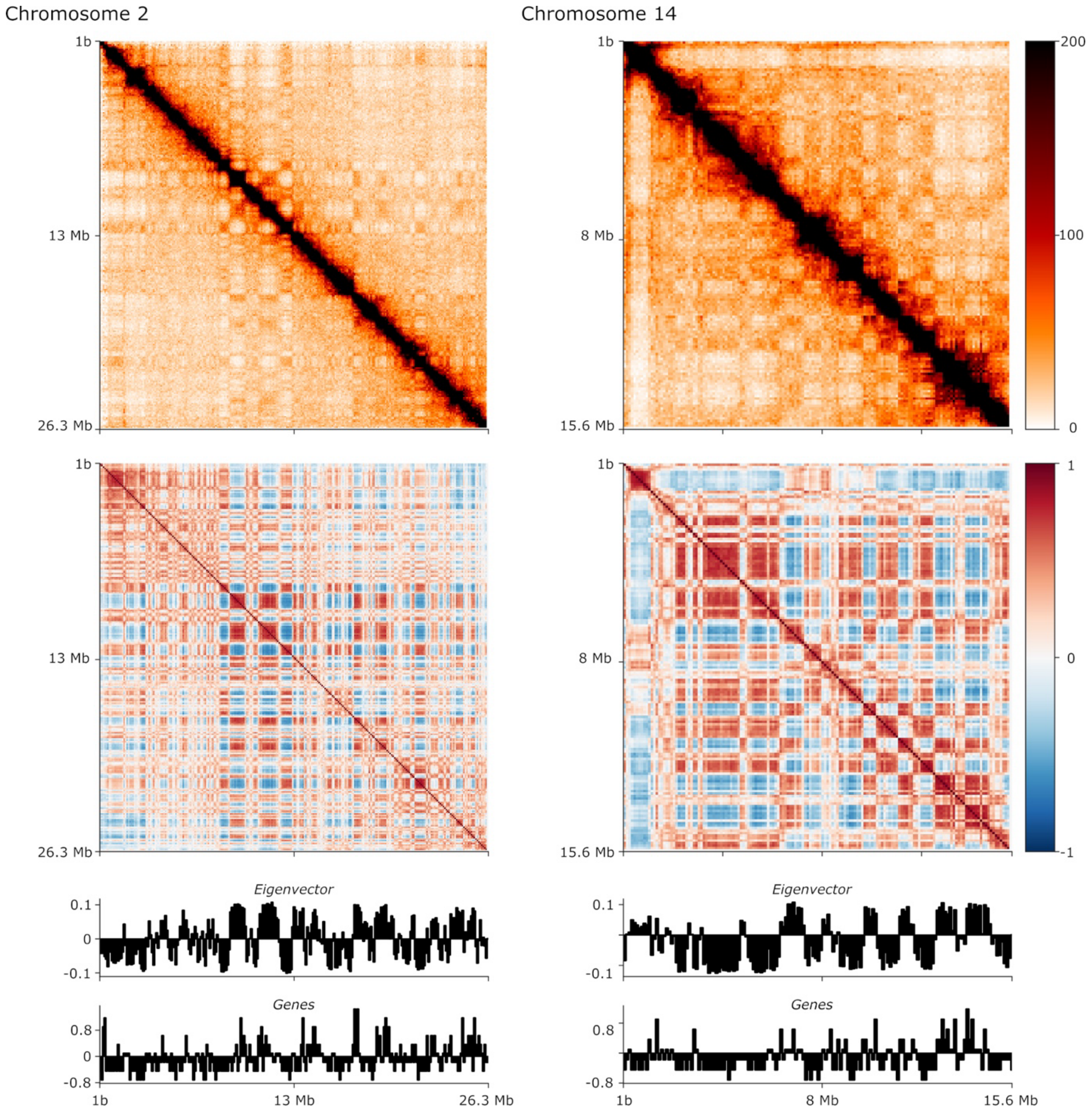


***Supplementary Figure 19. A/B compartment analyses in cyclostomes.*** *Top, Hi-C map from sea lamprey (Petromyzon marinus) liver. Middle, Pearson matrices showing A/B compartmentalization of chromosomes. Bottom, eigenvector and gene density plots (resolution = 100 kb). Note the enrichment in gene density for A compartment (positive eigenvector signal). Representative examples from chromosomes 2 and 14. Similar results are obtained in other chromosomes.*

**A**

1

$$\frac{\text{PbetaTub85D} - \text{FLP}}{\text{PbetaTub85D} - \text{FLP}};;\text{ry} \quad \textbf{X} \quad \frac{\text{FM7c (B, YFP)}}{-};;\frac{\text{TM2 (Ubx, e)}}{\text{TM6c (YFP, Tb, e)}}$$

2

$$\frac{\text{PbetaTub85D} - \text{FLP}}{\text{FM7c (B, YFP)}};;\frac{\text{ry}}{\text{TM6c (YFP, Tb, e)}} \quad \textbf{X} \quad \frac{\text{FM7c (B, YFP)}}{-};;\frac{\text{TM2 (Ubx, e)}}{\text{TM6c (YFP, Tb, e)}}$$

3

$$\frac{\text{PbetaTub85D} - \text{FLP}}{\text{FM7c (B, YFP)}};;\frac{\text{TM2 (Ubx, e)}}{\text{TM6c (YFP, Tb, e)}} \quad \textbf{X} \quad \frac{\text{PbetaTub85D} - \text{FLP}}{-};;\frac{\text{TM2 (Ubx, e)}}{\text{TM6c (YFP, Tb, [illegible]}}$$

$$\frac{\text{PbetaTub85D} - \text{FLP}}{\text{PbetaTub85D} - \text{FLP}};;\frac{\text{TM2 (Ubx, e)}}{\text{TM6c (YFP, Tb, e)}}$$

**B**

1

$$\frac{\text{PbetaTub85D} - \text{FLP}}{\text{PbetaTub85D} - \text{FLP}};;\frac{\text{TM2 (Ubx, e)}}{\text{TM6c (YFP, Tb, e)}} \quad \textbf{X} \quad \text{w[*]};;\frac{\text{3xFLAG FRT CTCF} - \text{GFP FRT mmCTCF}}{\text{3xFLAG FRT CTCF} - \text{GFP FRT mmCTCF}}$$

2

$$\text{w[*]};;\frac{\text{3xFLAG FRT CTCF} - \text{GFP FRT mmCTCF}}{\text{3xFLAG FRT CTCF} - \text{GFP FRT mmCTCF}} \quad \textbf{X} \quad \frac{\text{PbetaTub85D} - \text{FLP}}{-};;\frac{\text{3xFLAG FRT CTCF} - \text{GFP FRT mmCTCF}}{\text{TM6c (YFP, Tb, e)}}$$

3

$$\frac{\text{w[*]}}{\text{PbetaTub85D} - \text{FLP}};;\frac{\text{3xFLAG FRT CTCF} - \text{GFP FRT mmCTCF}}{\text{3xFLAG FRT CTCF} - \text{GFP FRT mmCTCF}} \quad \textbf{X} \quad \frac{\text{PbetaTub85D} - \text{FLP}}{-};;\frac{\text{3xFLAG FRT CTCF} - \text{GFP FRT mmCTC[illegible]}}{\text{TM6c (YFP, Tb, e)}}$$

$$\frac{\text{PbetaTub85D} - \text{FLP}}{\text{w[*]}};;\frac{\text{3xFLAG mmCTCF}}{\text{TM6c (YFP, Tb, e)}} \quad \textbf{OR} \quad \frac{\text{PbetaTub85D} - \text{FLP}}{\text{PbetaTub85D} - \text{FLP}};;\frac{\text{3xFLAG mmCTCF}}{\text{TM6c (YFP, Tb, e)}} \quad \textbf{OR}$$

$$\frac{\text{PbetaTub85D} - \text{FLP}}{-};;\frac{\text{3xFLAG mmCTCF}}{\text{TM6c (YFP, Tb, e)}} \quad \textbf{OR} \quad \frac{\text{w[*]}}{-};;\frac{\text{3xFLAG mmCTCF}}{\text{TM6c (YFP, Tb, e)}}$$

**C**

$$\frac{\text{PbetaTub85D} - \text{FLP}}{\text{w[*]}};;\frac{\text{3xFLAG mmCTCF}}{\text{3xFLAG mmCTCF}} \quad \textbf{OR} \quad \frac{\text{PbetaTub85D} - \text{FLP}}{\text{PbetaTub85D} - \text{FLP}};;\frac{\text{3xFLAG mmCTCF}}{\text{3xFLAG mmCTCF}} \quad \textbf{OR}$$

$$\frac{\text{PbetaTub85D} - \text{FLP}}{-};;\frac{\text{3xFLAG mmCTCF}}{\text{3xFLAG mmCTCF}} \quad \textbf{OR} \quad \frac{\text{w[*]}}{-};;\frac{\text{3xFLAG mmCTCF}}{\text{3xFLAG mmCTCF}}$$

***Supplementary Figure 20. Overview of fly crosses performed to obtain ortholog replacement lines. A.*** *Crosses performed to generate the fly line expressing FLP (on chr2) and double-balanced on chr3. The stable stock genotype is highlighted in the yellow box.* ***B.*** *Crosses performed to introduce FLP in lines carrying the ortholog downstream of the endogenous locus. Once GFP cannot be detected, the recombination has occurred, and only the exogenous ortholog is present. The stable stock genotypes are highlighted in the green box.* ***C.*** *All possible genotypes for the homozygous replacement flies. mmCTCF = mouse CTCF; a representative for any other ortholog.*

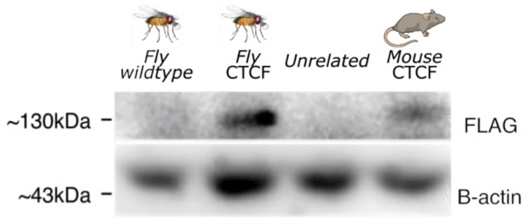


***Supplementary Figure 21. Western blot for fly replacement lines.*** *Original image from the Western blot displayed in Figure 3B, showing the unrelated lane that was removed for visualization purposes.*
